# Disruption of a CCR5-like immunoglobulin gene is linked to plague susceptibility in black-footed ferrets

**DOI:** 10.64898/2026.06.26.734856

**Authors:** Yana Safonova, Taylor Pursell, Caleb S. Whitley, Katelyn R. Sheneman, Anna Mikhailova, Vinamratha Pattar, Mariia Pospelova, Adonis A. Rubio, Katalin A. Voss, Jordan M. Welker, Anton Zamyatin, Anton Bankevich, Jef D. Boeke, Emily Haraguchi, Elizabeth Hudson, Eric Kline, Tanya M. Lama, William Lauer, Valerie Le Sage, Milton Thomas, Corey T. Watson, Shirong Zheng, Christopher O. Barnes, Seema S. Lakdawala, Matt Pennell, Melissa L. Smith, Scott Boyd, Matthew B. Lawrenz, Klaus-Peter Koepfli

**Author notes:** Corresponding authors (Y.S.), (S.B.), (M.B.L.), (K.-P.K.). These authors contributed equally to this work (Yana Safonova, Taylor Pursell, Caleb S. Whitley, Katelyn R. Sheneman).

## Abstract

Black-footed ferrets (*Mustela nigripes*) are among the world’s most endangered mammals and remain highly vulnerable to sylvatic plague caused by *Yersinia pestis*, yet the genetic basis of this susceptibility has remained unknown. Current conservation strategies rely on vaccination of captive-bred animals and large-scale flea control with insecticides, approaches that are costly, labor-intensive, and difficult to implement across the species’ natural range. Several closely related mustelid species, including the domestic ferret, are substantially more resistant to plague, providing an opportunity to identify naturally evolved immune mechanisms through comparative immunogenomics. Here we identify a conserved class of immunoglobulin lambda variable genes encoding unusually long antigen-binding loops with CCR5-mimicking sequence features that are widespread among Caniformia species. Because CCR5 has been implicated in host interactions with *Yersinia* species, we hypothesized that antibodies encoded by these germline genes contribute to plague resistance through receptor-like molecular mimicry. Consistent with this hypothesis, we show that these genes are under strong purifying selection in mustelids, are actively expressed in antibody repertoires, and monoclonal antibodies encoded by them reduced intracellular *Y. pestis* survival in macrophages. In contrast, all analyzed black-footed ferrets carried a frameshifting deletion resulting in loss of gene expression. These findings identify a naturally disrupted germline antibody gene as a candidate determinant of plague susceptibility in black-footed ferrets, demonstrating that variation in germline immunoglobulin genes can influence susceptibility to a lethal infectious disease. Ultimately, these findings lay the groundwork for genetically informed conservation management and the development of new antibody-based anti-plague strategies.

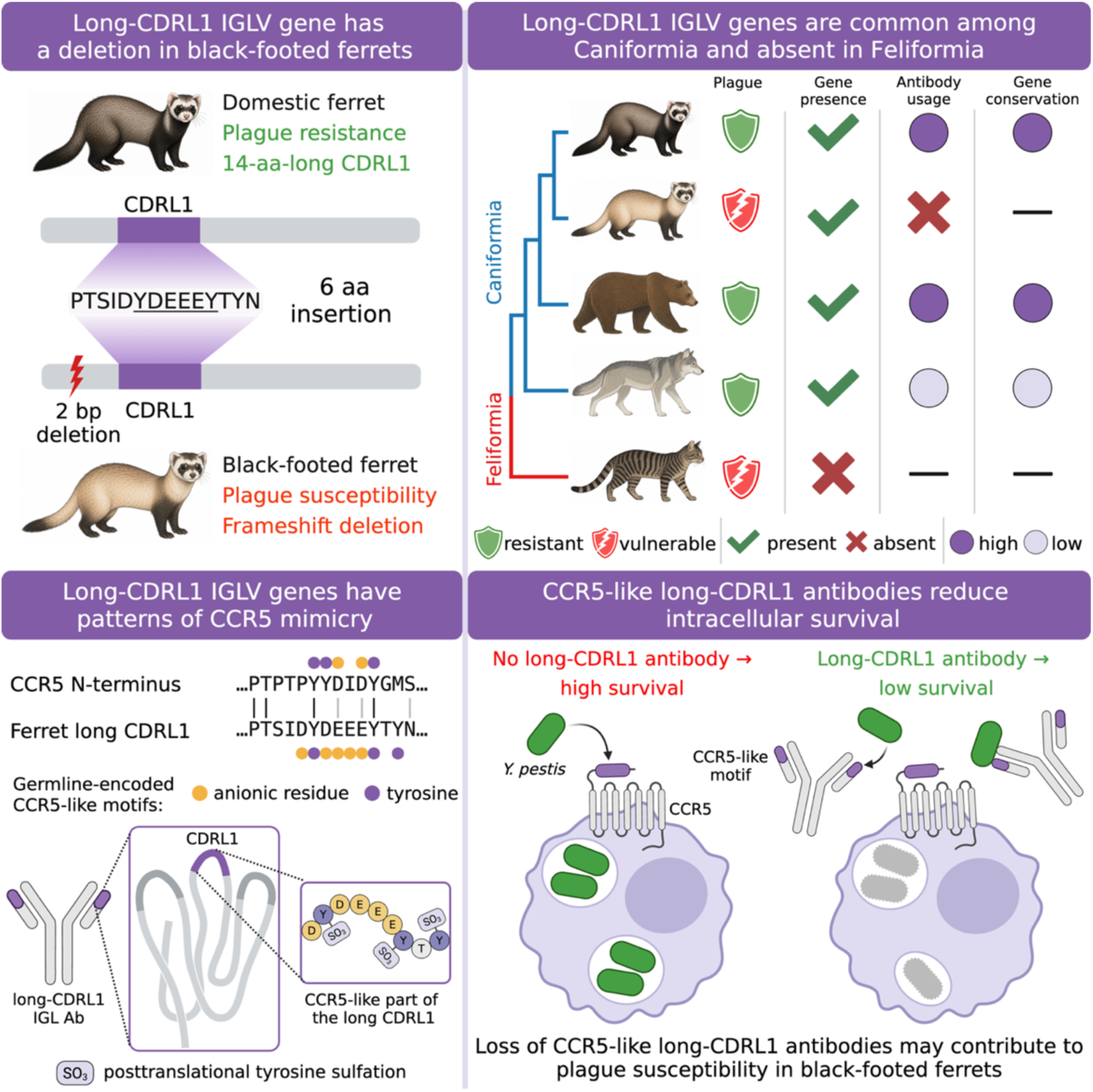

## Introduction

The black-footed ferret (*Mustela nigripes*) represents one of the most dramatic examples of population collapse and recovery in modern conservation biology. Historically distributed throughout the Great Plains of North America, black-footed ferrets depended on prairie dog colonies for both prey and shelter. During the early twentieth century, extensive eradication campaigns of prairie dogs, the main prey of black-footed ferrets, coupled with habitat destruction and infectious disease outbreaks caused a catastrophic decline of black-footed ferret populations (Jachowski, 2014). By the late 1970s, the species was presumed extinct until a small remnant population was rediscovered near Meeteetse, Wyoming, in 1981. However, this population rapidly collapsed following outbreaks of sylvatic plague and canine distemper, leaving only 18 animals by 1985 (Jachowski, 2014). To prevent extinction, the remaining ferrets were brought into captivity to establish a breeding program, yet only seven individuals successfully reproduced and founded the entire extant population. As a result, all living black-footed ferrets descend from these seven founders and retain less than 10% of the historical genetic diversity of the species (Wisely et al., 2002).

Despite intensive conservation efforts, sylvatic plague caused by *Yersinia pestis* remains one of the major threats to black-footed ferret recovery. The disease is transmitted to ferrets through the bites of infected fleas (primarily *Oropsylla hirsuta*) found on prairie dogs and other rodents or by consumption of infected prey (Mize et al., 2017). Captive-bred individuals are routinely vaccinated against plague prior to release, but vaccine protection is incomplete and difficult to scale across wild prairie dog ecosystems (Abbott et al., 2012). Interestingly, susceptibility to plague differs substantially among closely related mustelid species. Phylogenomic analyses demonstrated that black-footed ferrets are most closely related to the steppe polecat (*Mustela eversmanni*) of Central Asia and the European polecat (*Mustela putorius*), the progenitor of the domestic ferret (*Mustela putorius furo*) (Totikov et al., 2026). Experimental infection studies showed that domestic ferrets exhibit relative resistance to *Y. pestis* infection and develop strong antibody responses following bacterial challenge, whereas steppe polecats are substantially more susceptible (Williams et al., 1991). The contrasting plague phenotypes observed among closely related species suggest that variation in immune gene content or function may contribute to differential susceptibility within the Mustelinae subfamily. This pattern is similarly reflected across the mammalian order Carnivora, with species in some families showing high susceptibility to plague (cats, Felidae), while others are relatively resistant (foxes and wolves, Canidae; bears, Ursidae) (Gage & Kosoy, 2005; Oyston and Williamson, 2011).

To investigate these differences, germline immunoglobulin (IG) loci were selected as the basis for comparative analysis. IG loci encode antibodies, central mediators of adaptive immunity that recognize diverse pathogens and antigens, through somatic mechanisms including V(D)J recombination, junctional diversification, and somatic hypermutation (Hozumi and Tonegawa, 1976; Bernard et al., 1978; Alt and Baltimore, 1982). In most mammals, antibodies are encoded by three IG loci corresponding to the heavy and light chains of the antibody molecule: IGH (immunoglobulin heavy chain locus), IGK (immunoglobulin kappa light chain locus), and IGL (immunoglobulin lambda light chain locus). Each locus contains arrays of variable (V), diversity (D, IGH only), and joining (J) genes denoted as IGH/K/LV, IGHD, and IGH/K/LJ which recombine to form V(D)J sequences. The V(D)J sequences correspond to the variable regions of antibodies and contain three complementarity-determining regions (CDRs) that represent the antigen-binding site. These regions are referred to as CDRH1–3 for the heavy chain and CDRL1–3 for the light chain.

Although antibody diversity is often viewed primarily as a product of somatic diversification, growing evidence indicates that variation within germline IG genes can directly shape a composition of expressed V(D)J sequences and may be associated with responses to pathogens and inflammatory disorders (Avnir et al., 2016; Parks et al., 2017; Johnson et al., 2021; Pennell et al., 2023; Rodriguez et al., 2023; Engelbrecht et al., 2025). Comparative immunogenomics studies have characterized IG loci across mammals and demonstrated that they are among the most evolutionarily dynamic regions of the genome, often harboring extensive structural variation, lineage-specific expansions, and rapid gene speciation (Ota and Nei, 1994; Rodriguez et al., 2023; Pospelova et al., 2025; Yoo et al., 2025; Peres et al., 2026). However, despite the increasing number of IG gene annotations, the functional significance of germline diversity remains poorly understood. In particular, it remains unclear whether unusual germline-encoded antigen-binding features represent neutral evolutionary variation or adaptive solutions shaped by recurrent host-pathogen interactions.

In this work, we employed an integrated approach combining comparative genomics, antibody repertoire sequencing, and functional experiments across Carnivora species to identify and characterize a conserved immunoglobulin lambda chain variable (IGLV) gene with unusual evolutionary and functional properties that is potentially associated with plague susceptibility in mustelids. Specifically, our objectives were: 1) to map the evolutionary conservation and divergence patterns of the IGLV gene of interest across Carnivora, with particular attention to lineage-specific variations in mustelids; 2) to examine the sequence and structural-level features of antibodies encoded by the identified IGLV gene utilizing both computational predictions and experimental data; and 3) to propose a mechanistic link between the identified genetic IGLV variants and differential susceptibility to plague in mustelids, thereby elucidating the potential host factors underlying the variable penetrance of the disease. Our findings provide insights into how immune gene evolution shapes pathogen-host dynamics.

## Results

### Black-footed ferrets have a deletion in the IGLV gene with unusually long CDRL1

Unlike domestic ferrets, black-footed ferrets are highly susceptible to the sylvatic plague caused by the gram-negative, coccobacillus bacterium, *Yersinia pestis*. To reveal immune markers possibly associated with susceptibility to the plague, genomes of the domestic ferret and the black-footed ferret were sequenced using PacBio High Fidelity (HiFi) reads with 20–30x coverage and assembled using hifiasm (Cheng et al., 2021; see **Methods**). IG loci were annotated in the assembled genomes using the IgDetective tool (Sirupurapu et al., 2022), manually curated, and compared (see **Methods**). Comparison revealed an IGLV gene characterized by unusually long CDRL1: 42 nt compared to 27 nt in the closest human IGLV gene (IGLV1-40) and to ∼21 nt in other ferret IGLV genes according to the IMGT (the International ImMunoGeneTics information system; Lefranc, 2003) definition of CDRs (**Figure 1A, Figure S1**). In the Kabat definition (Kabat and Wu, 1971), the CDRL1 is extended by nine nt to the 5’ end and six nt to the 3’ end with a total length of 57 nt (**Figure 1A**). Because both IMGT and Kabat CDR definitions cover the insertion in the ferret IGLV gene of interest, they can be used interchangeably, and the IMGT definition was chosen as the default in all downstream analyses.

**Figure 1.**
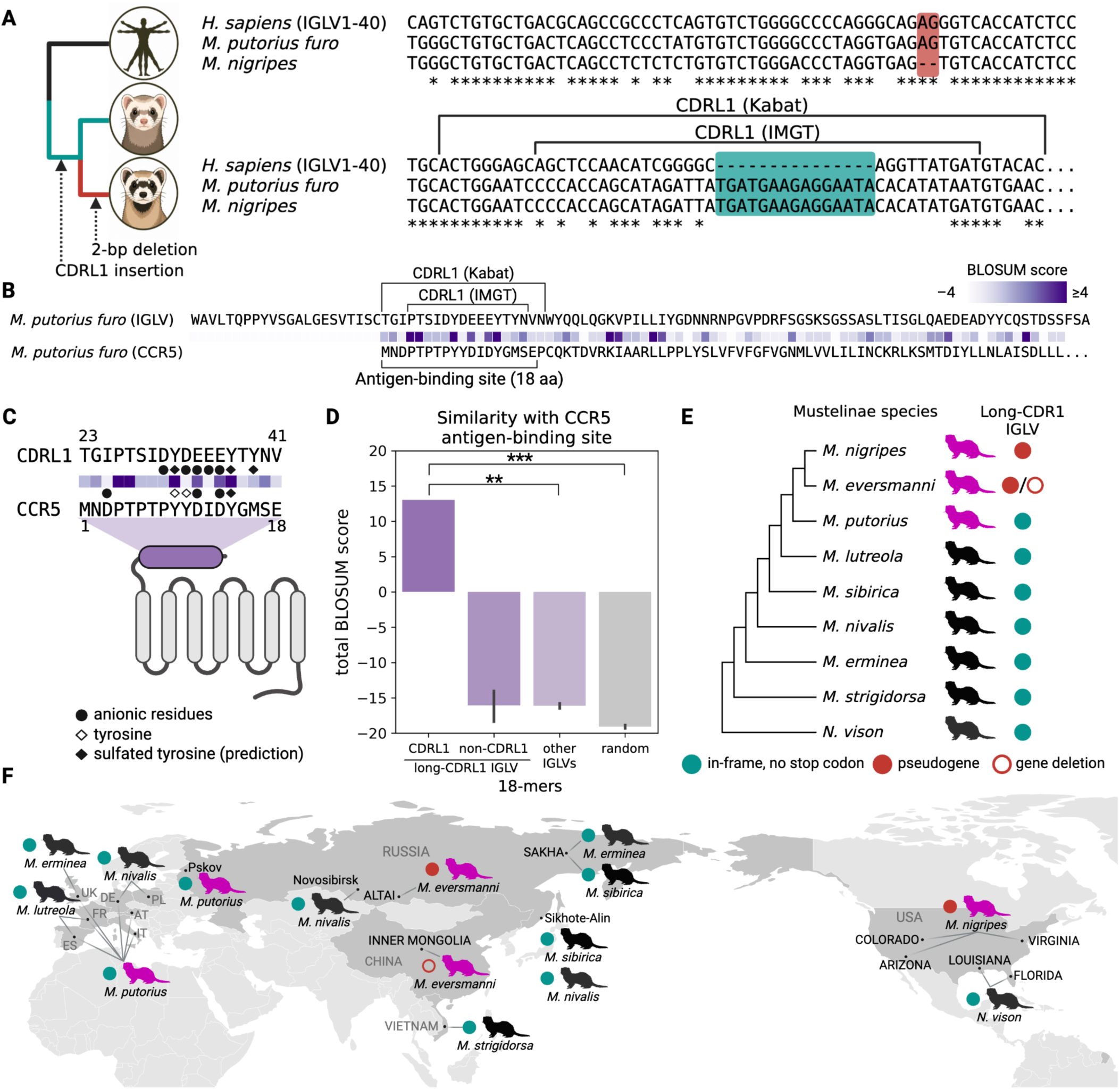
The landscape of long-CDRL1 IGLV genes across Mustelinae species. **(A)** Alignment of the 123-nucleotide long prefix of the CDRL1 IGLV gene in the domestic ferret (*Mustela putorius furo*, middle) against the prefixes of the closest human IGLV gene (IGLV1-40; top) and the orthologous gene in the black-footed ferret (*M. nigripes*, bottom). Positions of CDRL1 are shown by brackets; the CDRL1 insertion is shown by the green box; the two-nucleotide deletion in the black-footed ferret is shown by the red box. The complete alignment of three genes is shown in **Figure S1. (B)** Amino acid alignments of the long-CDRL1 IGLV gene (top line) against the prefix of CCR5 gene in the domestic ferret. Positions of predicted antigen-binding sites are shown by brackets. BLOSUM scores between corresponding amino acids are color coded by shades of violet: from –4 (pale) to ≥4 (dark). **(C)** Alignment of the 18-amino-acid long fragment of the IGLV gene corresponding to the start of CDRL1 (Kabat notation) and the antigen-binding site in CCR5 starting from the N-terminus end. BLOSUM scores are shown in shades of violet (see the legend of panel B); special features of amino acid sequences are highlighted by black circles (anionic residues), hollow diamonds (tyrosines), and black diamonds (sulfated tyrosines predicted by Sulfinator). **(D)** Total BLOSUM scores computed between the 18-aa long binding site of the domestic ferret CCR5 and (from left to right) the 18-aa long CDRL1 fragment shown in panel B; all other 18-mers of the target IGLV gene; 18-mers of all other IGLV genes; randomly generated 18-mers. Here and further, bars show 95% confidence intervals. Empirical P-values are encoded as follows: ns:P>0.05, *<0.05, **<0.01, ***<0.001, ****<0.0001. **(E)** Mustelinae species selected for the analysis. The cladogram is based on the phylogenetic tree in Totikov et al., 2026. Members of the subgenus Putorius are shown by magenta. The green filled circles indicate in-frame long-CDRL1 IGLV genes without stop codons, and red circles show non-productive long-CDRL1 IGLV genes. The hollow red circle indicates that no long-CDRL1 IGLV genes were detected. **(F)** Map showing the geographic locations of Mustelinae samples and productivity of the long-CDRL1 IGLV gene. Countries corresponding to shown locations are shown in darker gray color. The rest of the legend is consistent with panel E.

The long-CDRL1 IGLV gene in the domestic ferret has an in-frame sequence and does not contain stop codons, whereas its ortholog in the black-footed ferret genome has a two-nucleotide deletion leading to a shift in the open reading frame and likely resulting in a loss of function (**Figure 1A**, **Figure S1**).

### Long-CDRL1 IGLV gene has CCR5-mimicking features

**Figure 1B** shows that 9 out of 14 amino acids in the detected long CDRL1 are either anionic (aspartate, D; glutamate, E) or tyrosines (Y). Anionic motifs as well as posttranslation sulfation of tyrosines were reported as key features of antigen-binding loops of antibodies neutralizing HIV by mimicking the surface protein CCR5 that is used by the virus to invade host cells (Huang et al., 2004). These motifs are not encoded in the antigen-binding loops of germline human IG genes and only emerge in CDRH3s as the result of random insertions and deletions introduced during V(D)J recombination (Choe et al., 2003; Huang et al., 2004). Previous studies suggested that CCR5-mediated interactions may influence host susceptibility during *Yersinia* infection and contribute to macrophage targeting by *Y. pseudotuberculosis*, the progenitor of *Y. pestis* (Sheahan and Isberg, 2015). Although the molecular mechanisms underlying these interactions remain incompletely understood, the unusual CCR5-like features of the long-CDRL1 IGLV gene raise the possibility that antibodies derived from it may possess receptor-like binding properties relevant to *Yersinia*-host interactions and contribute to early protective immune responses against infection. Consistent with this hypothesis, the two-nucleotide deletion in the black-footed ferret’s long-CDRL1 IGLV gene likely renders it non-functional.

To analyze the similarity between CCR5 and the long-CDRL1 IGLV gene in more detail, the domestic ferret CCR5 gene was extracted using alignment of the human CCR5 gene via minimap2 (Li, 2018). Because previous studies showed that antigens recognize the sulfated N-terminus of CCR5 spanning the first ∼18 amino acids (Huang et al., 2007), the 18-amino-acid long prefix of the domestic ferret CCR5 gene was used as approximation of the CCR5 antigen-binding site and referred to as *CCR5 18-mer*. The long CDRL1 possesses the features of the CCR5 18-mer: abundant anionic residues and CDRL1 tyrosines that are aligned to CCR5 tyrosines and predicted to be sulfated by the Sulfinator tool (Monigatti et al., 2002) (**Figure 1C, 1D**). To confirm similarity, the CCR5 18-mer was aligned to the long-CDRL1 IGLV gene using local alignment, and the score was computed as the sum of BLOSUM scores (Henikoff & Henikoff, 1992) between pairs of aligned amino acids. The fragment covering the CDRL1 has the best alignment score: 13 versus an average of –16 for other 18-mers in the same IGLV gene (**Figure 1E**). This score also exceeds scores obtained from aligning the CCR5 18-mer to other domestic ferret IGLV genes (average score –16; empirical P=0.002) and to randomly generated amino acid 18-mers (N=5000, average score –19, empirical P=0.0006) (**Figure 1F**).

### Long-CDRL1 IGLV genes are found in most Mustelinae species

To investigate the genetic landscape of long-CDRL1 IGLV genes among the closest relatives of the domestic ferret, we collected whole-genome sequencing (WGS) data, comprising both genome assemblies and short-read sequences, from species within the Mustelinae subfamily. In total, 30 subjects of nine species were selected: the Western polecat (*Mustela putorius*, 7 subjects), the steppe polecat (*Mustela eversmanni*, 2 subjects), the black-footed ferret (8 subjects); the back-striped weasel (*Mustela strigidorsa*, 1 subject); the Siberian weasel (*Mustela sibirica*, 2 subjects), the European mink (*Mustela lutreola*, 2 subjects), the least weasel (*Mustela nivalis*, 4 subjects), the stoat (*Mustela erminea*, 2 subjects), and the American mink (*Neogale vison*, 2 subjects) (**Table S1**). The geographical locations of the selected subjects are spread over the European and Asian parts of Eurasia as well as North America.

The domestic ferret long-CDRL1 IGLV gene was mapped using minimap2 (Li, 2018) to each WGS dataset and identified in 29 out of 30 subjects (**Figure 1D**). All eight black-footed ferrets share the frameshift-causing two-nucleotide deletion in the long-CDRL1 IGLV gene identified in the genome assembly, suggesting that black-footed ferrets are likely homozygous at this locus (**Figure 1E**). The same deletion was also found in the genome of the Altai steppe polecat (**Figure 1E**). Because black-footed ferrets and steppe polecats are sister species (Totikov et al., 2026), this observation suggests the deletion emerged before the two species diverged. Another steppe polecat (Inner Mongolia, China) was the only subject without matches to the long-CDRL1 IGLV gene (**Figure 1E**). Because the Altai steppe polecat has an available genome assembly (GCA_963422785.1), reads of the Inner Mongolia steppe polecat were aligned to it. **Figure S2** shows that the genomic position of the long-CDRL1 IGLV gene coincides with reduced read coverage in the Inner Mongolia steppe polecat. This might be explained by either sequencing artefacts or structural variants in the steppe polecat population. For the other 20 subjects representing seven species, in-frame sequences without stop codons were identified for the long-CDRL1 IGLV gene (**Figure 1E**). The fact that the in-frame long-CDRL1 IGLV gene and its defective ortholog were found across both Eurasia and North America supports that the two-nucleotide deletion is explained by phylogeny rather than differences in antigens on continents (**Figure 1E**).

### Long-CDRL1 IGLV genes are common among dog-like carnivoran species

To determine whether the long-CDRL1 IGLV gene emerged earlier in the order Carnivora, we conducted comparative analysis using publicly available genome assemblies from 40 carnivoran species alongside the sequenced genomes of the domestic ferret and the black-footed ferret (**Figure 2A, Table S2**). Additionally, genomes of the cheetah (*Acinonyx jubatus*), the gray wolf (*Canis lupus lupus*), the maned wolf (*Chrysocyon brachyurus*), the red panda (*Ailurus fulgens*), and the sloth bear (*Melursus ursinus*) were sequenced using Pacbio HiFi to ∼20x coverage and assembled using hifiasm (Cheng et al., 2021). To adequately resolve the complex architecture of IGL loci, only genomes with a contig N50 above 200 kbp were used. Information about samples and genome assembly characteristics are summarized in **Tables S3**, **S4**. In total, 33 Caniformia (dog-like carnivores) and 14 Feliformia (cat-like carnivores) representing 10 taxonomic families were analyzed (**Figure 2A**). All genomes were processed using IgDetective (Sirupurapu et al., 2022), and detected IGLV genes were aligned using IgBlast (Ye et al., 2013) to identify CDRL1s. **Figure 2A** shows that 30 out of 33 Caniformia species have in-frame long-CDRL1 IGLV genes without stop codons: for 24 of them, the CDRL1s were 14 aa long, and, for the remaining six species, the CDRL1s were 13 aa long and differed from the 14-aa-long CDRL1s by one-amino-acid deletion. For canids and bears, up to three copies of the long-CDRL1 IGLV gene were found. In addition to the black-footed ferrets, an out-of-frame long-CDRL1 IGLV gene was found for the Northern sea otter (*Enhydra lutris kenyoni*). It contains a different one-nucleotide deletion that shifts the reading frame and results in a premature stop codon (**Figure S3**). Long-CDRL1 IGLV genes were not detected in the plains spotted skunk (*Spilogale interrupta*), which may be an artifact of locus fragmentation, as the IGL locus is split across three distinct contigs in the genome assembly. Unlike Caniformia species, all Feliformia species appear to lack long-CDRL1 IGLV genes; the longest CDRL1s are 9 aa long and are found across all Feliformia IGLV genes.

**Figure 2.**
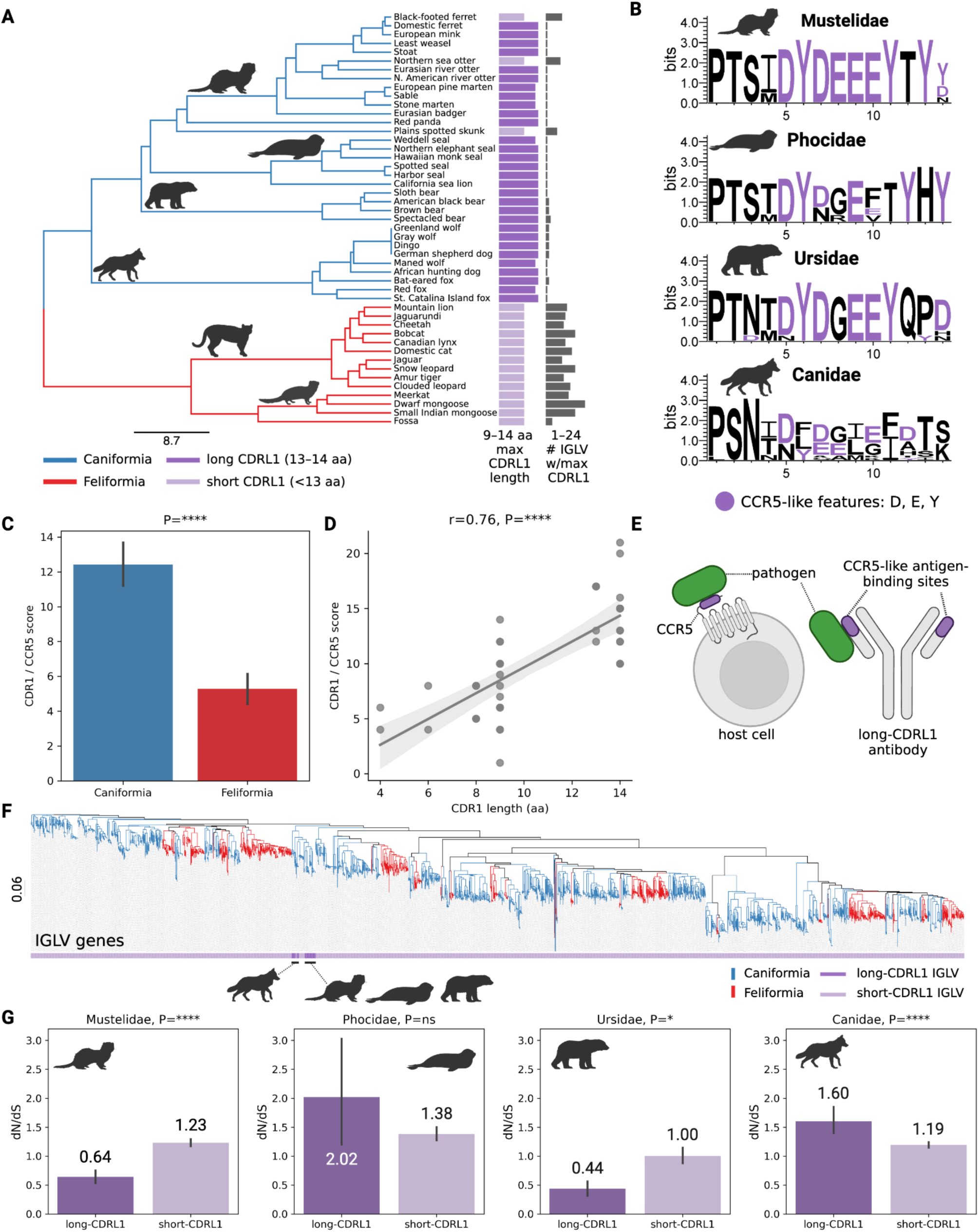
Long-CDRL1 IGLV genes across Carnivora species. **(A)** The phylogenetic tree of Carnivora species selected for comparative genomic analysis. The tree was pruned from the VertLife mammalian tree (https://vertlife.org/data/mammals; Upham et al., 2019) and, for subspecies of *Canis lupus*, a polytomy at the terminal branch corresponding to this lineage was introduced. Caniformia and Feliformia species are shown by blue and red branches, respectively. Branches corresponding to taxonomic families with more than one species (Mustelidae, Phocidae, Ursidae, Canidae, Felidae, Herpestidae) are marked with species silhouettes. Bars on the right show the length of the longest CDRL1 across in-frame IGLV genes without stop codons of the corresponding species (bright violet if 13-14 aa long; pale violet if shorter than 13 aa) and the number of IGLV genes with CDRL1s of this length. Here and further, phylogenetic trees are visualized using Iroki (Moore et al., 2020). **(B)** Weblogos (Crooks et al., 2004) of amino acid sequences of long CDRL1s found in species from the Mustelidae, Phocidae, Ursidae, and Canidae families. Anionic residues and tyrosines are shown in violet. **(C)** CCR5 / CDRL1 scores in Caniformia and Feliformia suborders. Here and further, P-values were computed using the Kruskal-Wallis test unless otherwise specified. **(D)** CCR5 / CDRL1 scores versus lengths of corresponding CDRL1s across all species in panel A. The Pearson’s correlation and corresponding P-value are shown at the top. **(E)** A proposed mechanism of molecular CCR5 mimicry in long-CDRL1 IGLV antibodies. **(F)** The phylogenetic tree of in-frame IGLV genes without stop codons collected across all species shown in panel A and computed using Clustal Omega (Sievers et al., 2011). Blue and red branches correspond to IGLV genes of Caniformia and Feliformia species, respectively. Bright and light violet colors in the bar below the tree correspond to long- and short-CDRL1 IGLV genes, respectively. **(G)** The distributions of dN/dS values computed for long- and short-CDRL1 IGLV genes for all pairs of species in the Mustelidae, Phocidae, Ursidae, and Canidae families. Numbers show average values of the distributions.

### Long CDRL1s can be diverse but retain CCR5-like features

Sequences of 13–14 aa long CDRL1s vary across Caniformia families: while the known signature of CCR5 mimicry described above (abundant tyrosines and anionic residues) is strong in mustelids and ursids, it is less pronounced in phocids and canids (**Figure 2B**). To account for divergence of CCR5 sequences across species, CCR5 sequences were extracted from the genomes and the IGLV gene with the closest CDRL1 match was found for each species. **Figure 2C** shows that the CCR5 / CDRL1 similarity scores computed as the sum of BLOSUM scores positively correlate with the CDRL1 lengths (Pearson’s correlation r=0.76, P=7.65×10^−10^). The CCR5 / CDRL1 similarity scores are higher in the Caniformia species compared to the Feliformia species (P=5.37×10^−7^, the Kruskal-Wallis test; **Figure 2D**). CCR5 / CDRL1 similarity scores across Caniformia families shows that even though the 14 aa long CDRL1s of canids do not carry the known CCR5 mimicking signature, they still resemble respective antigen-binding sites of canid CCR5s (**Figure S4**). Together these observations suggest that most Caniformia species have germline genes with a potential to generate CCR5-like IGL antibodies that possibly emerged due to a prolonged exposure to CCR5-interacting pathogens (**Figure 2E**).

### Long-CDRL1 IGLV genes are subjected to purifying selection in ferrets and bears

A phylogenetic tree of IGLV genes collected across the selected carnivoran species shows that long-CDRL1 IGLV genes are assorted into several clades: one encompasses long-CDRL1 IGLV genes of species comprising the infraorder Arctoidea (Ursidae, Phocidae, Otariidae, Mephitidae, Ailuridae, and Mustelidae) and three others consist of Canidae long-CDRL1 IGLV genes (infraorder Cynoidea) (**Figure 2F**). To understand differences in the evolutionary development of long-CDRL1 IGLV genes, analysis of non-synonymous to synonymous substitution rates (dN/dS) was performed for all IGLV genes for four Caniformia families that are present in more than one species (Canidae, Mustelidae, Phocidae, Ursidae). For each pair of species (*s*1, *s*_2_) from the same taxonomic family, the closest IGLV gene in *s*_2_ was computed for each IGLV gene in *s*_1_ and the dN/dS value was calculated. Species with non-productive long-CDRL1 IGLV genes were excluded from the analysis, and only in-frame IGLV genes without stop-codons were used. **Figure 2G** shows the distributions of dN/dS values for long-CDRL1 IGLV genes versus short-CDRL1 IGLV genes for four taxonomic families. In Mustelidae and Ursidae, long-CDRL1 IGLV genes have dN/dS values <1 that are lower compared to short-CDRL1 IGLV genes, suggesting that they are under strong purifying selection (Mustelidae: P=5.41×10^−7^; Ursidae: P=0.02; Kruskal-Wallis test). Long-CDRL1 IGLV genes of Phocidae species have dN/dS values >1 that are not different from short-CDRL1 IGLV genes. In Canidae species, long-CDRL1 IGLV genes have dN/dS values >1 that are also higher compared to short-CDRL1 IGLV genes, indicating positive selection (P=1.91×10^−8^; Kruskal-Wallis test). Across all families, short-CDRL1 IGLV genes exhibit average dN/dS values close to one, suggesting that most of them are evolving neutrally.

### Germline variation in IGLV genes shapes expressed repertoires

While the results of genomic analyses suggest that long-CDRL1 IGLV genes with CCR5-mimicking signatures are subjected to high selective pressure in Mustelidae and Ursidae (**Figure 2G**), their functionality in expressed antibody repertoires has been undetermined. To test whether long-CDRL1 IGLV genes contribute to functional repertoires, RNA-seq libraries containing expressed antibody repertoires were generated for six Caniformia species: domestic ferret (Mustelidae), black-footed ferret (Mustelidae), gray wolf (Canidae), maned wolf (Canidae), red panda (Ailuridae), and sloth bear (Ursidae). For the black-footed ferret, gray wolf, maned wolf, red panda, and sloth bear, PacBio Iso-Seq data were generated from spleen samples of the same animals used for whole genome sequencing (**Figure 3A**; see **Methods**). For the domestic ferret, repertoire-sequencing (Rep-Seq) libraries were generated from paired PBMCs and spleen using the 5’ RACE and IGLC primers for three new animals (**Figure 3A**; see **Methods**). Information about read counts for generated Rep-Seq and Iso-Seq libraries are summarized in **Table S5, S6**. Additionally, publicly available Rep-Seq data generated from PBMCs of domestic dogs (*Canis lupus familiaris*) (Cullen et al., 2022) and domestic ferrets (Hebert et al., 2025) were added to the dataset (**Figure 3A**; **Table S7**). While a mixture of tissues and technologies is a potential source of batch effects, downstream analyses demonstrated consistent results across all datasets.

**Figure 3.**
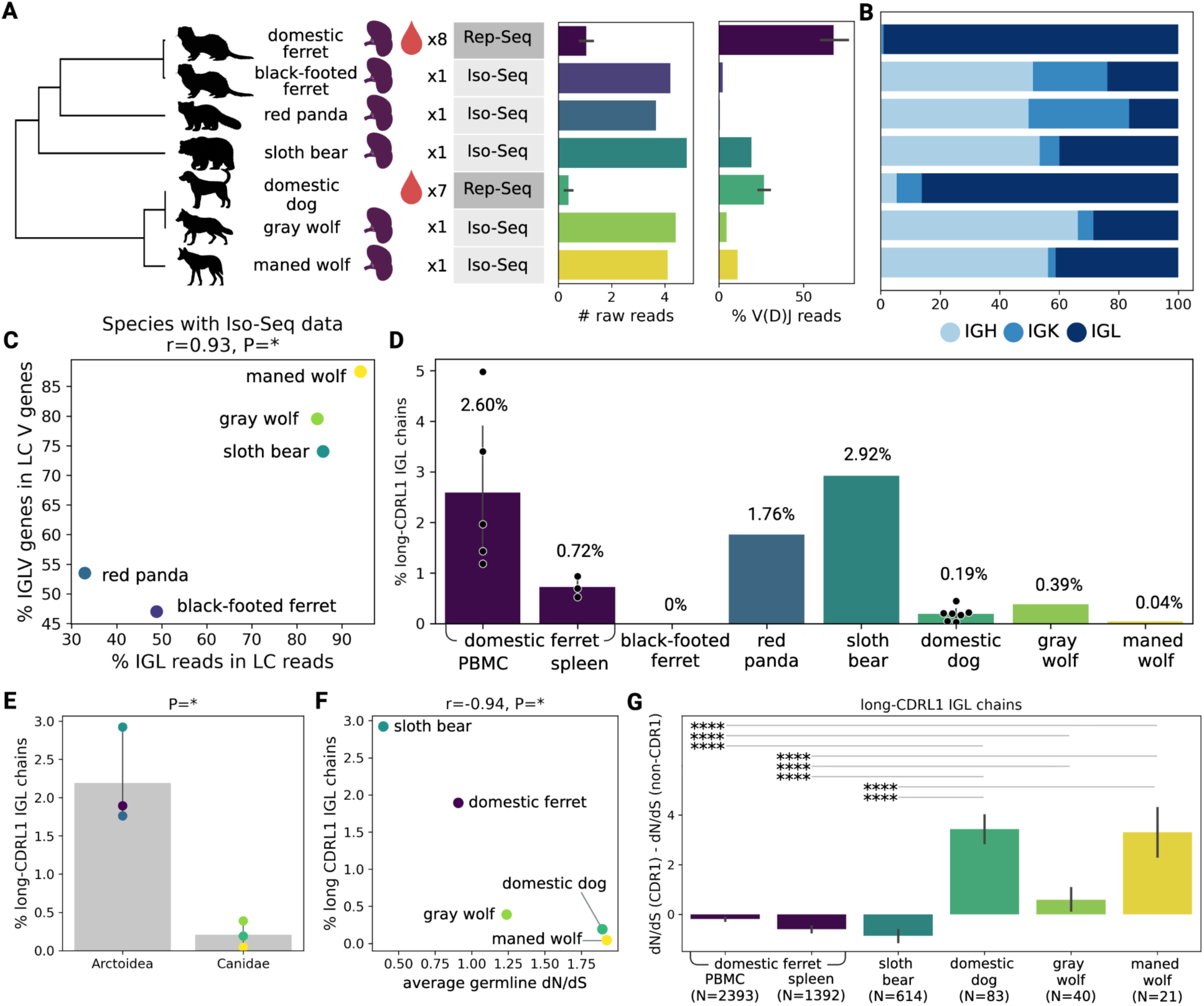
Analysis of expressed long-CDRL1 IGLV across carnivores. **(A)** Description of expressed antibody repertoire sequencing data used for analysis including a list of species, tissues (spleen or PBMC), the number of samples, and the technology (Rep-Seq or Pacbio Iso-Seq). Left and right barplots show the counts of raw reads in a library and percentages of V(D)J sequences, respectively. **(B)** The distribution of IGH, IGK, and IGL V(D)J sequences across all analyzed species. For species with more than one library (the domestic ferret, the domestic dog), average values were computed. **(C)** Percentage of IGL reads in all light chain reads versus percentage of IGLV genes in all light chain V genes across five species with Iso-Seq data (black-footed ferret, red panda, sloth bear, gray wolf, maned wolf). **(D)** Percentages of long-CDRL1 IGL sequences across analyzed samples. **(E)** Percentages of long-CDRL1 IGL sequences in Arctoidea species versus Canidae species. Only species with productive long-CDRL1 IGLV genes were selected. **(F)** Average long-CDRL1 IGLV gene dN/dS values in Figure 2G versus average percentages of long-CDRL1 sequences in expressed IGL repertoires across five species: sloth bear, domestic ferret, gray wolf, domestic dog, and maned wolf. **(G)** Differences between dN/dS of CDRL1 and dN/dS of the remaining parts of the V gene across long-CDRL1 IGL sequences of five carnivore species. The counts of sequences used for the analysis are shown on the bottom.

To analyze expressed antibody repertoires using RNA-sequencing data shown in **Figure 3A**, annotated IG loci from genomes of corresponding species were transformed into germline databases and integrated into the DiversityAnalyzer tool (Shlemov et al., 2017). RNA-sequencing libraries were aligned to the constructed germline databases and reads corresponding to V(D)J recombinations were recruited and classified according to the IG chain type: IGH, IGK, or IGL. While IGL chains are dominant in targeted Rep-Seq data for the domestic ferret and the domestic dog by design, Iso-Seq data likely contain proportions of IG chains that are more identical to ones in the natural repertoires. **Figure 3B** shows that the fraction of IGL sequences is higher in Iso-Seq data of the sloth bear, gray wolf, and maned wolf. The percentages of IGL sequences among all light chain sequences correlates positively with the percentages of IGLV genes among all light chain V gene sequences (Pearson’s correlation r=0.93, P=0.02; **Figure 3C**). An anticorrelation between the counts of IGKV and IGLV genes was previously observed across mammalian species (Pospelova et al., 2025). The correlation in **Figure 3C** suggests that proportions of IGLV and IGKV genes are directly translated into proportions of IGK and IGL transcripts.

### Conserved long-CDRL1 IGLV genes have higher usages in expressed antibody repertoires

The IGL sequences with long CDRL1s were detected in expressed antibody repertoires of all analyzed species except for the black-footed ferret. The latter is consistent with the frameshift-causing two-nucleotide deletion found in all analyzed black-footed ferrets (**Figure 1D, 1E**). Alignment of recombined IGL sequences with long CDRL1s revealed all such IGLs are derived from the long-CDRL1 IGLV genes and do not result from somatic hypermutations (SHMs). The percentage of unique long-CDRL1 IGL sequences computed with respect to the total number of unique IGL sequences varies from species to species. The highest percentages were detected in Arctoidea species: the sloth bear (2.92%), the domestic ferret (1.89%; averaged across all samples), and the red panda (1.76%) (**Figure 3D**). In the domestic ferrets, the percentage of long-CDRL1 IGL sequences is higher in PBMCs compared to the spleen (P=0.025; Kruskal-Wallis test). The lowest percentages of long-CDRL1 IGL sequences were detected in Canidae species: gray wolf (0.39%), domestic dog (0.19%; averaged across all samples), and maned wolf (0.04%) (**Figure 3D**). The difference in percentages between the Arctoidea and Canidae groups is statistically significant (P=0.049; Kruskal-Wallis test; **Figure 3E**). The percentages of long-CDRL1 IGL sequences also anticorrelate with average dN/dS values of germline long-CDRL1 IGLV genes computed in comparison with species from the same family as shown in **Figure 2A** and **2G** (Pearson’s correlation r=–0.94, P=0.02; **Figure 3F**). This observation suggests that more conserved IG genes have higher usage in expressed antibody repertoires.

Long-CDRL1 IGLV genes have different levels of SHMs across analyzed species (**Figure S5**), which is likely linked to the differences in their usages. To compare SHM signals in these genes across species, dN/dS levels were computed for CDRL1s and non-CDRL1 parts of the V gene. The red panda was excluded from the analysis as it only had 19 long-CDRL1 IGL sequences. While dN/dS levels of CDRL1s are lower in domestic ferrets, the differences between species can be influenced by individual immune responses and are thus difficult to interpret (**Figure S6**). However, the distribution of differences between the dN/dS values for CDRL1s and non-CDRL1s within the same sequence computed across all long-CDRL1 IGL sequences shows a clear pattern: the differences are negative for the domestic ferret and sloth bear sequences and are lower compared to the domestic dog, gray wolf, and maned wolf that are characterized by positive differences (**Figure 3G**). This suggests that the CDRL1s of conserved long-CDRL1 IGLV genes shared across Arctoidea tolerate fewer non-synonymous SHMs than those of the more diverged IGLV genes found among Canidae species. Analysis of expanded clonal lineages would help to understand the role of SHMs in long-CDRL1 IGL sequences.

### Long-CDRL1 IGL antibodies in domestic ferrets have unique signatures

To analyze properties of long-CDRL1 chains in antibodies, single-cell RNA sequencing (scRNA-Seq) data of expressed antibody repertoires of domestic ferrets were collected and analyzed. First, two scRNA-seq libraries were generated from PBMCs of a new ferret subject (denoted as subject S1) (**Figure 4A**; see Methods). Second, publicly available scRNA-Seq libraries corresponding to the spleen of the second animal (subject S2, 4 replicas) and PBMCs of the third animal (subject S3, 4 replicas) were used (**Figure 4A**) (Walsh et al., 2025). For each animal, scRNA-Seq libraries were assembled into full-length transcripts using MiXCR (Bolotin et al., 2015) or 10x Genomics Cell Ranger (v10.0.0) and annotated using DiversityAnalyzer (Shlemov et al., 2017). Only cell barcodes containing one heavy chain and one light chain sequence were selected for further analysis. The number of selected antibodies varied across subjects: S1 (601–614), S2 (1236–1294), S3 (1729–1812), with the percentage of IGL antibodies ranging from 43.09% to 55.47% (**Figure 4A, Table S8**). The percentage of long-CDRL1 IGL antibodies in all IGL antibodies ranged from 3.28% to 5.02% (**Figure 4A, Table S8**), which is similar to the percentages of long-CDRL1 IGL chains found in unpaired PBMC antibody repertoires and is higher compared to unpaired spleen antibody repertoires (**Figure 3D**).

**Figure 4.**
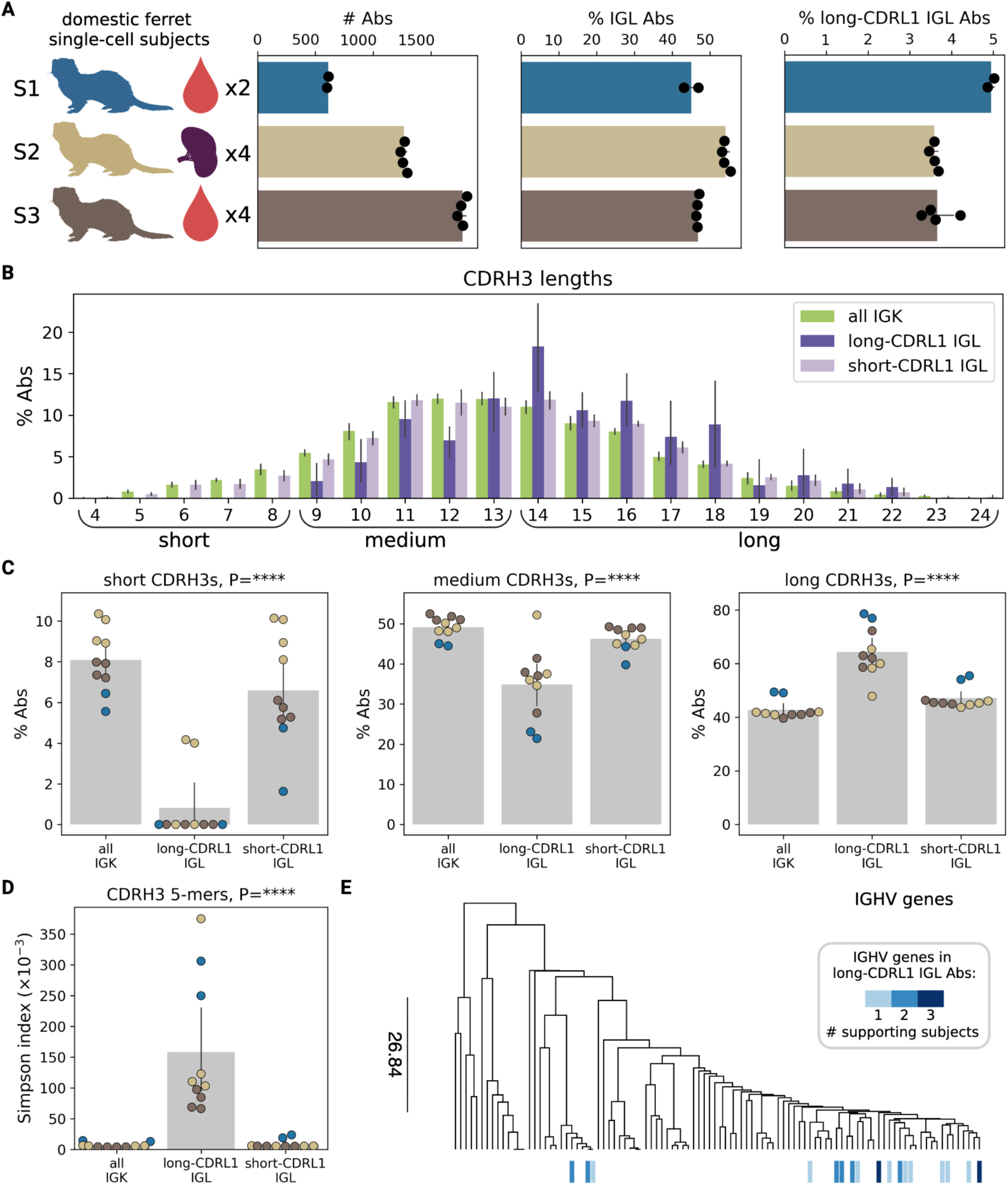
Long-CDRL1 antibodies in single-cell data of domestic ferrets. **(A)** Description of single-cell RNA-seq data of expressed antibody repertoires of three domestic ferrets (S1, S2, S3) including tissues (spleen or PBMC) and the number of replicas. Barplots show (from left to right): counts of antibodies with one heavy and one light chain; percentages of IGL antibodies in all antibodies; percentages of long-CDRL1 IGL antibodies in all IGL antibodies. **(B)** The distribution of CDRH3 lengths (in aa) across long-CDRL1 IGL Abs (violet), short-CDRL1 Abs (light violet), and IGK Abs (green). **(C)** The cumulative percentages of Abs with short, medium, and long CDRH3s across long-CDRL1 IGL Abs, short-CDRL1 IGL antibodies, and IGK Abs. **(D)** Values of the Simpson index computed using frequencies of CDRH3 amino acid 5-mers with a multiplicity 2 or more across long-CDRL1 IGL Abs, short-CDRL1 IGL antibodies, and IGK Abs. **(E)** The guide tree of IGHV genes computed using hierarchical clustering. IGHV genes used in long-CDRL1 IGL Abs are marked by blue bars according to the number of supporting subjects: 1 (light blue), 2 (medium blue), and 3 (dark blue).

Long-CDRL1 IGL antibodies are characterized by longer CDRH3 sequences compared to short-CDRL1 IGL antibodies and IGK antibodies (**Figure 4B**). While short CDRH3s (≤8 aa) are present in ∼6.59% of short-CDRL1 IGL antibodies and ∼8.08% of IGK antibodies, they are rare in long-CDRL1 IGL antibodies (0.82% on average) (P=1.40×10^−21^, the linear mixed-effects model; **Figure 4C**). CDRH3s with medium lengths (9–13 aa) have slightly lower percentages in long-CDRL1 IGL antibodies: ∼34.89% compared to ∼46.29% in short-CDRL1 IGL antibodies and ∼49.12% in IGK antibodies, contributing to a statistically significant difference (P=3.11×10^−9^, the linear mixed-effects model; **Figure 4C**). Long CDRH3s (≥14 aa) contribute to ∼64.29%, ∼47.11%, and ∼42.80% of long-CDRL1 IGL, short-IGL antibodies, and IGK antibodies, respectively (P=9.09×10^−20^, the linear mixed-effects model; **Figure 4C**).

To estimate the sequence diversity of CDRH3s of long-CDRL1 IGL antibodies, their amino acid 5-mers were extracted, and frequencies were computed as fractions of CDRH3s containing them. Simpson indices were then computed as the sum of squared frequencies across all scRNA-Seq libraries. Similar computations were performed for short-CDRL1 IGL antibodies and IGK antibodies. To account for the smaller numbers of long-CDRL1 IGL antibodies that can result in higher frequencies of rare 5-mers, only 5-mers with a multiplicity 2 or more were considered. **Figure 4D** shows higher Simpson index values in the long-CDRL1 IGL antibodies, suggesting a lower diversity of their CDRH3 sequences (P=7.17×10^−10^, the linear mixed-effects model). In particular, one 5-mer (DSYGY) representing middle parts of CDRH3s and two 5-mers (DYLDV, YAMGY) corresponding to CDRH3 suffixes have significantly higher fractions in long-CDRL1 IGL antibodies, indicating preferential usages of one ferret IGHD and two IGHJ genes (**Figure S7**). Finally, 17 IGHV genes with higher usage in long-CDRL1 IGL antibodies were detected, however, only two IGHV genes have higher usages in all three subjects (**Figure 4E**). This can be explained by sampling effects as well as germline differences between subjects S1–S3 associated with IGHV gene usages. **Figure 4E** shows that 14 out of 17 IGHV genes are closely related, suggesting that preferences toward usage of specific IGHV genes in long-CDRL1 IGL antibodies are still possible.

### Long-CDRL1 IGL antibodies have sulfotyrosines

Sulfation of N-terminal tyrosines of CCR5 are critical to its function. These sulfotyrosines not only enhance native ligand affinity (Seibert et al., 2002; Abayev et al., 2018) but are also required for interactions with HIV-1 gp120 enabling viral entry (Farzan et al., 1999; Jen et al., 2009). As Tyrosylprotein Sulfotransferase 1 and 2 (TPST1 and TPST2) do not have a single, rigid consensus sequence, validation of sulfation requires *in vitro* expression. To validate the sulfation of CDRL1-containing light chains, five native pairs were selected from the single cell data to capture sequence diversity while preserving the conserved CDRL1 motif. Pairs were chosen to represent: (i) the most prevalent IGH/IGL pairings, (ii) three clones (FTP2, 3, and 4) sharing an IGL sequence but divergent IGH chains, isolating effect of heavy-chain sequences on sulfation; (iii) variation in CDR3L length across the three different IGL sequences to probe whether adjacent loop length influences sulfation; and (iv) diversity of CDR3H length and sequence. Sequences were humanized by placing the V(D)J ferret sequences into a human IgG1/IgL backbone (**Figure 5A**).

**Figure 5.**
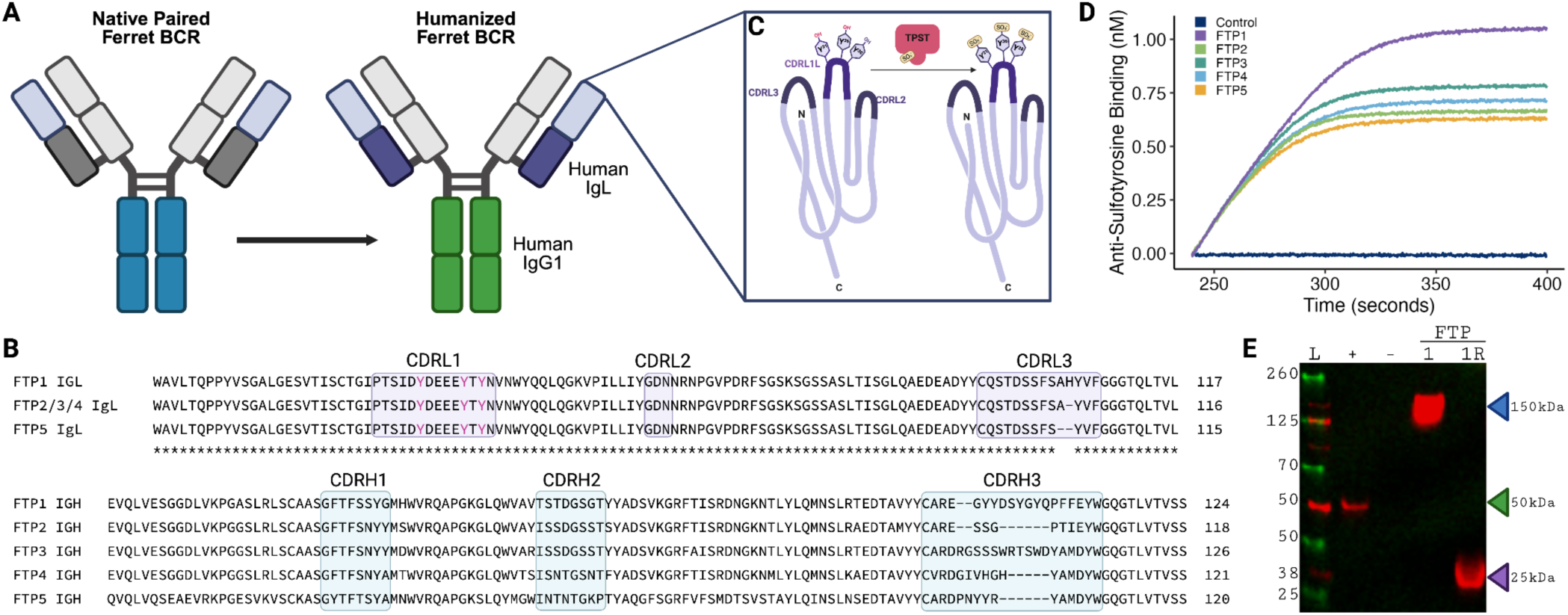
Patterns of tyrosine sulfation in long-CDRL1 IGL mAbs. **(A)** Schematic showing conversion of native paired ferret B cell receptors (BCR) to humanized antibody format with human IgG1 and IgL constant regions. **(B)** Sequence alignment of selected ferret antibody clones (FTP1-FTP5) heavy and light chain sequences highlighting CDR1, CDR2, and CDR3 regions and potentially sulfated tyrosines. **(C)** Structured schematic of antibody tyrosine sulfation in the CDRL1 by TPST. **(D)** Line plot of binding kinetics of anti-sulfotyrosine monoclonal to humanized ferret antibodies (FTP1-FTP5) and control (mAb 515) measured by biolayer interferometry (BLI). **(E)** Western blot of monoclonals either intact or reduced (R) probed with anti-suflotyrosine. Molecular weight of intact monoclonal, heavy chain, light chain, and Fab indicated.

The selected ferret clones (FTP1-FTP5) maintained the conserved CDRL1 (**Figure 5B**) but with unique CDRL3 sequences. Three native pairs (FTP2, 3, and 4) share the same IGL sequence but have diverse CDRH3 lengths in IGH chains. Human and ferret TPST2 share 84% amino acid identity and all four catalytic residues (R78, E99, K158 and S285) as well as the 5’-phosphoribosyl-1-pyrophosphate synthase (5’PSB) motif (GVPRSGTTL) are conserved between both species; we therefore concluded that ferret TPST2 should behave similarly to that from human. The monoclonals were co-expressed with human TPST2 and screened for sulfation with biolayer interferometry (BLI) (**Figure 5C**). Although all clones showed sulfation (**Figure 5D**) compared to a non-sulfated, human control monoclonal, we observed varying signal intensities with the highest value for FTP1. The differences in BLI saturation levels across clones can be attributed to variable sulfation stoichiometry, heterogeneous sulfation patterns across multiple tyrosine residues within the CDRL1 regions (**Figure 5C**), and clone-specific differences in local sequence context that influence anti-sulfotyrosine antibody binding affinity. To confirm the sulfation is on the light chain, a western blot comparing intact versus reduced clones was run. Unlike the non-sulfated control, all clones demonstrated anti-sulfotyrosine reactivity with the intact antibody (∼150 kDa) (**Figure 5E**), confirming our BLI findings. Under reducing conditions, anti-sulfotyrosine reactivity at ∼25 kDa was observed while no signal was detected at ∼50 kDa, consistent with the sulfation only occurring on the light chain (**Figure 5E**). Together, these data demonstrate that light chain sulfation occurs during recombinant expression and could occur *in vivo*, thus enhancing the CCR5-mimicry potential.

### Long-CDRL1 IGL antibodies decrease intracellular survival of *Yersinia pestis*

CCR5 has been implicated as a potential receptor for *Y. pestis* that facilitates interactions with leukocytes that benefit the bacteria (Sheahan and Isberg, 2015). Therefore, if long-CDRL1 antibodies containing CCR5-like domains act as a CCR5 mimic, it is possible that the antibodies may alter interactions between *Y. pestis* and macrophages. To test this hypothesis, *Y. pestis* was grown for 3 hours at 37°C prior to macrophage infection to induce expression of the Ysc type 3 secretion system (T3SS) and the Yop effectors that are secreted via this system. As expected, growth at 37°C and expression of the T3SS limited the ability of macrophages to phagocytose the bacteria as compared to *Y. pestis* grown at 26°C (**Figure 6A**). While opsonization with naïve human serum increased phagocytosis of *Y. pestis* grown at 37°C (**Figure 6A**, P=0.03, the one-way ANOVA test), it did not increase the ability of the macrophages to kill the phagocytosed bacteria (**Figure 6B**). Similarly, opsonization with serum from two domestic ferrets, F99-22 and F103-22, significantly increased phagocytosis (**Figure 6A**, P<0.0001; the one-way ANOVA test). However, unlike human serum, opsonization with domestic ferret serum significantly limited bacterial survival within macrophages (**Figure 6B**, P=0.004 and P<0.0001 for F99-22 and F103-22, respectively; the one-way ANOVA test). This result suggests that factors in domestic ferret serum, but absent in human serum, alter the ability of *Y. pestis* to survive within macrophages. To specifically test if long-CDRL1 antibodies directly contribute to this phenotype, *Y. pestis* was incubated with the natively paired long-CDRL1 humanized domestic ferret mAbs (FTP1-FTP5) described above, or the murine mAb 7.3, which recognizes the T3SS-associated LcrV protein to inhibit Yop effector translocation through the T3SS and promotes phagocytosis (Ivanov et al., 2014). While mAb 7.3 incubation increased phagocytosis, the FTP antibodies did not significantly increase phagocytosis (**Figure 6C**). However, the most sulfated monoclonal antibody, mAb FTP1 (**Figure 5D**), significantly decreased *Y. pestis* intracellular survival (**Figure 6D**, P=0.04; the one-way ANOVA test) in comparison to the other sulfated mAbs and murine anti-LcrV mAb 7.3 (**Figure 6D**). In support of this finding, correlation analysis suggests sulfation levels influence the function of the long-CDRL1 antibodies (**Figure 6E**, Pearson’s correlation r=–0.95, P=0.02). Together these data indicate that long-CDRL1 antibodies can influence *Y. pestis* interactions with macrophages, limiting the ability of the bacteria to evade macrophage killing.

**Figure 6.**
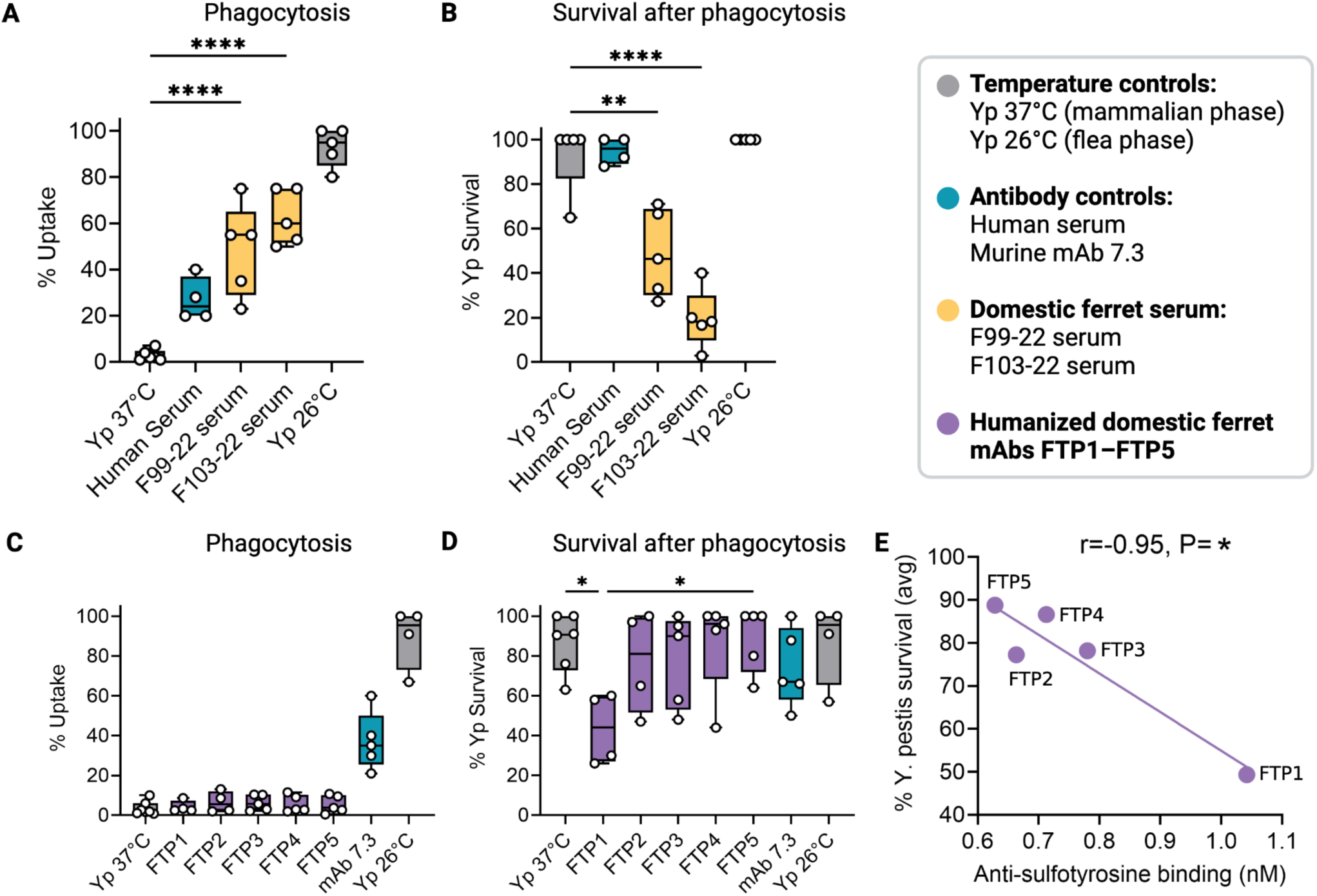
Opsonization of *Y. pestis* with domestic ferret serum or sulfated mAb improves bacterial killing by macrophages. RAW 264.7 macrophages were infected with *Y. pestis* KIM1001 grown at 37°C opsonized with either (**A, B**) domestic ferret serum or (**C, D**) sulfated mAbs (FTP1-FTP5). (**A**) Percent of phagocytosed bacteria and (**B**) intracellular bacteria recovered after 6 hours when opsonized with 5% serum. (**C**) Percent of phagocytosed bacteria and (**D**) intracellular bacteria recovered after 6 hours when opsonized with sulfated mAbs FTP1-FTP5 (100 µg/mL). Non-opsonized bacteria grown at 26°C or 37°C or opsonization with human serum or murine anti-LcrV mAb 7.3 included as controls. (**E**) Comparison of anti-sulfotyrosine binding and mean intracellular *Y. pestis* survival within macrophages for monoclonal antibodies FTP1-FTP5. Statistical significance was determined by the one-way ANOVA with Tukey’s multiple comparison test in panels A–D and Pearson’s correlation analysis in panel E.

## Discussion

### Long-CDRL1 IGLV genes in mustelids

The black-footed ferret (*Mustela nigripes*) is an endangered species native to North America and highly susceptible to sylvatic plague caused by *Yersinia pestis* (Godbey et al., 2006; Matchett et al. 2010). In contrast, closely related domestic ferrets (*Mustela putorius furo*), which descend from Western polecats (*M. putorius*) in Eurasia, demonstrate substantial resistance to plague and do not develop severe disease (Williams et al., 1991). While plague is endemic to both Eurasia and the Americas, research suggests that it has existed in Eurasia for at least 6,000 years (Achtman et al., 1999; Sun et al., 2014; Rasmussen et al., 2015; Rascovan et al., 2019), whereas it was introduced to North America only at the beginning of the 20th Century (Biggins and Kosoy, 2001; Barbieri et al. 2020). The close phylogenetic relationship between these species, juxtaposed with their distinct plague exposure histories during evolution, provides an opportunity to identify immunogenomic differences potentially associated with divergent responses to infection. In this study, comparative analyses identified a previously uncharacterized IGLV gene containing an unusually long CDRL1 (14 amino acids according to the IMGT definition in Lefranc, 2003) with sequence features resembling the antigen-binding region of the host chemokine receptor CCR5 in domestic ferrets. In black-footed ferrets, this IGLV gene carries a two-nucleotide deletion that disrupts the open reading frame and corrupts the predicted amino acid sequence. These observations led to the hypothesis that antibodies derived from this gene in domestic ferrets may exhibit CCR5-like molecular mimicry, whereas black-footed ferrets may have lost this potential functionality. Although the relationship between plague susceptibility and CCR5-like interactions remains unclear, we suggest that this locus may contribute to differences in immune responses to *Y. pestis*.

Molecular mimicry represents an additional mechanism by which antibodies can evolve receptor- or ligand-like binding solutions to neutralize pathogens (Cunningham, 2009). Examples include anti-idiotypic antibodies that structurally resemble antigens (Fields et al., 1995), receptor-mimicking antibodies targeting snake toxins (Ducancel et al., 1996), and influenza antibodies that mimic sialic-acid receptor interactions (Lee et al., 2014; Schmidt et al., 2015). HIV-1 broadly neutralizing antibodies that mimic the CCR5 N-terminus and act as CCR5 decoys during recognition of the viral envelope glycoprotein represent another example of molecular mimicry that is particularly relevant to this study (Choe et al., 2003; Dorfman et al., 2006; Huang et al., 2007; Walker et al., 2009). Structural and biochemical studies demonstrated that these interactions depend on sulfated tyrosine residues and acidic features within the antigen-binding site (Choe et al., 2003; Huang et al., 2004; Huang et al., 2007; Acharya et al., 2011). Notably, the long-CDRL1 IGLV gene identified in domestic ferrets contains multiple tyrosines aligned with those of CCR5, together with an enrichment of anionic residues, suggesting the potential for analogous receptor-like interactions.

### Long-CDRL1 IGLV genes in carnivores

The discovery of this long-CDRL1 IGLV gene raised broader questions: how widespread such genes are across carnivorans, whether they contribute to expressed antibody repertoires, and whether their CCR5-like biochemical features can be experimentally validated. To address these questions, a comparative immunogenomics study integrating evolutionary analyses, bulk and single-cell antibody repertoire sequencing, and biochemical characterization was performed. These analyses revealed that long-CDRL1 IGLV genes are common and evolutionarily conserved across dog-like carnivores and contribute to expressed antibody repertoires in multiple species. In domestic ferrets, antibodies derived from these genes not only retain CCR5-like sequence features but also contain posttranslationally sulfated tyrosines, supporting the possibility of receptor-like molecular mimicry. Together, these findings establish a link between the evolution of germline IG genes and functional antibody repertoires and suggest that germline-encoded features can preconfigure modes of antigen recognition.

At the genomic level, long-CDRL1 IGLV genes are broadly distributed across Caniformia species and absent in Feliformia, suggesting that they emerged early during the evolution of dog-like carnivorans. Their presence across mustelids, bears, canids, pinnipeds and other arctoid families indicates that these genes represent a conserved evolutionary innovation rather than lineage-specific anomalies. Notably, black-footed ferrets and steppe polecats (*Mustela eversmanni*) share the same two-nucleotide deletion disrupting the long-CDRL1 IGLV gene. A recent phylogenomic analysis of mustelids (Totikov et al., 2026) identified these species as sister taxa, which suggests that the deleterious mutation likely emerged in their common ancestor. Another Caniformia species carrying a deleterious mutation in the long-CDRL1 gene is the Northern sea otter (*Enhydra lutris kenyoni*). The deletion is one-nucleotide long and located at a different position than the deletion in black-footed ferrets and steppe polecats suggesting that these two deletions emerged independently from each other. All analyzed cat-like carnivorans possess substantially shorter CDRL1 regions and completely lack long-CDRL1 IGLV genes with CCR5-like features. This phylogenetic partitioning suggests that major carnivoran clades may differ in germline-encoded mechanisms of immune recognition and complements prior evidence for clade-associated differences in carnivoran immune phenotypes (Heinrich et al., 2016).

Evolutionary analyses further revealed lineage-specific selective pressures on long-CDRL1 IGLV genes. In Mustelidae and Ursidae, these genes are subject to strong purifying selection, suggesting functional constraints and conservation. In contrast, Canidae have signatures of positive selection consistent with diversification. These differences suggest that the functional role of long-CDRL1 antibodies may vary across carnivoran lineages and could reflect distinct pathogen pressures or ecological niches.

### Long-CDRL1 IGLV genes in antibody repertoires

To analyze links between germline variation and function, expressed antibody repertoires of six carnivoran species (black-footed ferret, domestic ferret, gray wolf, maned wolf, red panda, sloth bear) were sequenced and analyzed together with publicly available antibody repertoire data of domestic dogs. Long-CDRL1 IGLV genes are utilized in expressed repertoires in all species except for the black-footed ferret carrying a frameshifting deletion in this gene. These results confirm the differences in antibody repertoires of domestic ferrets and black-footed ferrets. The usage of long-CDRL1 IGLV genes correlates with their evolutionary conservation in other species. Species in which these genes are under purifying selection (ferrets and bears) show a higher usage in expressed repertoires compared to canid species with signatures of diversification. This inverse relationship between dN/dS and repertoire usage suggests that conserved genes may contribute to more functionally advantageous antibodies.

Single-cell analyses further showed that antibodies carrying long CDRL1s preferentially pair with heavy chains containing longer CDRH3s and biased IGHV, IGHD, and IGHJ gene usage patterns. These repertoire constraints suggest that long-CDRL1 IGL antibodies may participate in specialized recognition modes rather than broadly diverse antigen binding. Consistent with this interpretation, long-CDRL1 IGL antibodies contain biochemical features previously associated with CCR5-like interactions, including acidic residues and sulfated tyrosines. Biochemical experiments on five monoclonal long-CDRL1 IGL antibodies FTP1-FTP5 demonstrated that tyrosine residues within the light chain can undergo posttranslational sulfation. The detection of sulfotyrosines in these antibodies indicates that these residues are accessible to cellular sulfotransferases such as TPST1 and TPST2 and may contribute to binding interactions *in vivo*, supporting the hypothesis that these antibodies may adopt receptor-like binding properties.

### Possible CCR5 mimicry in responses to plague

Together, our findings suggest a previously undescribed form of receptor-like molecular mimicry encoded within the germline IGL locus. Although direct binding to CCR5 or CCR5 interactions with pathogens were not assessed in this study, the observed sequence features, evolutionary conservation, and repertoire usage patterns are consistent with the possibility that long-CDRL1 IGLV genes encode preconfigured solutions for interacting with CCR5-like targets. These observations further raise the possibility that long-CDRL1 IGLV genes may contribute to differential susceptibility to CCR5-interacting pathogens across carnivorans such as parasite *Toxoplasma gondii* (Ibrahim et al., 2010) as well bacteria *Staphylococcus aureus* (Alonzo et al., 2013) and *Yersinia pseudotuberculosis* (Sheahan and Isberg, 2015).

The latter is linked to the *Yersinia pestis*, a pathogen of particular interest acting as the causative agent of plague in both humans and mammalian wildlife (Barbieri et al., 2020; Mahmoudi et al., 2021). The distribution of long-CDRL1 IGLV genes is broadly consistent with reported differences in plague susceptibility among carnivoran species. Black-footed ferrets and steppe polecats both carry the out-of-frame long-CDRL1 IGLV gene and are highly vulnerable to plague (Williams et al., 1991). Domestic cats and several wild felids, including bobcats and mountain lions, lack long-CDRL1 IGLV genes and are known to be highly susceptible to *Y. pestis*, whereas dogs and other canids generally demonstrate greater resistance (Gage and Kosoy, 2005; Salkeld and Stapp, 2006; Oyston and Williamson, 2011). Serological studies further suggest that many dog-like carnivorans carrying long-CDRL1 IGLV genes, including bears, foxes, and raccoons, are frequently exposed to plague in endemic regions yet often survive infection and develop detectable antibody responses (Bevins et al., 2021).

Because long-CDRL1 IGLV genes likely emerged in the Caniformia lineage long before the estimated origin of *Y. pestis* ∼6,000 years ago (Achtman et al., 1999; Sun et al., 2014; Rasmussen et al., 2015; Rascovan et al., 2019), they were likely shaped by interactions with other pathogens. One possible candidate is *Yersinia pseudotuberculosis*, the evolutionary progenitor of *Y. pestis* (Rascovan et al., 2019), which split from *Yersinia enterocolitica* ∼0.4–1.9 millions of years ago (Achtman et al., 1999). Previous studies have implicated CCR5-associated host pathways in *Yersinia* infections. Tollenaere et al., 2008 reported associations between CCR5 polymorphisms and plague resistance in black rats (*Rattus rattus*) in Madagaskar, whereas Sheahan and Isberg, 2015 demonstrated that CCR5-associated host pathways influence *Y. pseudotuberculosis* infection, including reduced type 3 secretion effector translocation in CCR5-deficient macrophages. However, infection of CCR5-deficient BALB/c mice with an attenuated strain of *Y. pestis* suggests that complete loss of CCR5 may not significantly change plague susceptibility (Mecsas et al., 2004). More recent work suggests that the FPR1 receptor can also facilitate *Y. pestis* interactions with immune cells and may compensate for loss of CCR5 in the previous model (Osei-Owusu et al., 2019). Regardless of whether potential compensation mechanisms exist, CCR5 mimicry that would result in receptor-binding competition could affect plague pathogenesis differently than a loss-of-CCR5-function mutation. Together, these observations raise the possibility that CCR5-like mimicry may have evolved in response to exposure to Y. *pseudotuberculosis* or related ancestral pathogens and predisposed some carnivoran lineages to be more resistant to plague infection through a distinct mechanism.

Additionally, two species in which severe toxoplasmosis mortality has been extensively documented, black-footed ferrets (Burns et al., 2003) and sea otters (Miller et al., 2020), carry independently emerged deleterious mutations of the long-CDRL1 IGLV gene. While purely correlative, together these observations raise the possibility that loss of long-CDRL1 IGLV gene function may influence responses to certain pathogens.

### Long-CDRL1 IGL antibodies reduce intercellular survivability of *Y. pestis*

To explore whether long-CDRL1 IGL antibodies can influence host interactions with *Y. pestis*, we evaluated monoclonal antibodies FTP1-FTP5 in a mouse macrophage infection assay. Although none of the antibodies substantially altered bacterial uptake by macrophages, one monoclonal antibody (FTP1) significantly reduced intracellular *Y. pestis* survival following infection. This observation suggests that long-CDRL1 IGL antibodies can influence host-pathogen interactions during plague infection and that their activity may extend beyond classical opsonization-mediated enhancement of phagocytosis. Notably, the anti-plague activity of FTP1 was accompanied by the strongest anti-sulfotyrosine signal among the five tested monoclonal antibodies. Although this correlation is based on a limited number of antibodies and was largely driven by FTP1, it is consistent with the proposed model of CCR5-like molecular mimicry and supports the hypothesis that sulfotyrosine-associated features of long-CDRL1 IGL antibodies contribute to anti-plague activity.

### Future directions

Our study has several limitations. First, while sequence, biochemical, evolutionary, and functional observations are consistent with the proposed model of CCR5-like molecular mimicry, direct structural and mechanistic validation will be required to determine the precise targets and mode of action of long-CDRL1 IGL antibodies. Although one long-CDRL1 IGL monoclonal antibody demonstrated anti-plague activity, these observations do not establish a causal role for CCR5 mimicry. The proposed relationship between long-CDRL1 genes and plague susceptibility therefore requires further experimental validation, including direct binding studies, receptor competition assays, and infection models. Additionally, plague resistance in carnivorans is likely a polygenic trait shaped by multiple immune components and host-pathogen interactions, rather than by variation at a single immunoglobulin locus. Second, repertoire analyses were performed using a limited number of individuals and tissues and therefore may not fully capture the diversity of germline immunoglobulin genes and expressed antibody repertoires within and across populations.

Despite these limitations, our results establish a foundational framework for comparative immunogenomics to enable future studies of plague resistance and advance black-footed ferret conservation. They suggest that adaptive immune systems can encode preexisting solutions at the germline level, enabling rapid and efficient responses to recurrent challenges. Extending this framework to other rapidly evolving immune loci may reveal additional examples of germline-encoded specialization and clarify links between genome evolution, pathogen pressure, and immune function. In black-footed ferrets, these findings may also inform future conservation strategies aimed at restoring or introducing protective immune variation including targeted gene-editing approaches or other heritable immunogenomic interventions directed at plague susceptibility.

## Data availability

Raw sequencing data and genome assembled were deposited to NCBI. WGS and Iso-Seq data for black-footed ferret, cheetah, gray wolf, maned wolf, red panda, and sloth bear are available in BioProject PRJNA1472602 under accession SRR38916649–SRR38916659 [<u>reviewer link</u>]; corresponding genome assemblies are available via accessions PRJNA1472716–PRJNA1472727. Domestic ferret data is available in BioProject PRJNA1477079. Rep-Seq and single cell BCR libraries for domestic ferrets are available under accession numbers SRR39093867–SRR39093874 [<u>reviewer link</u>]. WGS for the domestic ferret are available via accession SRR39093875. Scripts and computed statistics are available at Github: <u>github.com/yana-safonova/CCR5-like_IGLV_genes</u>.

## Contributions

Y.S. made the initial observation and conceived the hypothesis linking the disrupted long-CDRL1 IGLV gene to plague susceptibility. Y.S., T.P., S.B., M.B.L., and K.-P.K. conceptualized the study. Y.S., T.P., E.Hu., E.K., W.L., M.L.S., M.T., C.T.W., S.Z., M.Pe., S.B., K.-P.K. generated the sequencing data. S.S.L. and V.L.S. provided domestic ferret samples. K.-P.K. provided the samples collected from the animals at the Smithsonian’s National Zoo and Conservation Biology Institute while he was a Research Associate. T.P., C.S.W., K.R.S., A.A.R., C.O.B., E.Ha., T.M.L., S.B., M.B.L. designed and performed wet-lab experiments. Y.S., T.P., C.S.W., K.R.S., A.M., V.P., M.Po., K.A.V., J.M.W., A.Z., A.B., J.D.B., M.B.L., M.Pe. analyzed the data. Y.S., T.P., C.S.W., V.P. prepared the figures. Y.S., T.P., C.S.W., M.B.L., K.- P.K. wrote the manuscript. All authors reviewed, commented on, and approved the manuscript.

## Acknowledgements

Y.S. was supported by the National Science Foundation under Grant No. 2536095. We are grateful to Raiees Andrabi, Ph.D., and Sergei Kliver, Ph.D., for invaluable discussions. We appreciate Dr. Pamela Bjorkman for sharing the TPST2 expression plasmid. We express our sincere gratitude to the dedicated animal care staff, veterinary technicians, veterinary pathologists, and researchers at the Smithsonian’s National Zoo and Conservation Biology Institute’s Center for Animal Care Sciences and Center for Species Survival, including Neel Aziz, D.V.M.; Adrienne Crosier, Ph.D.; Kristina Delaski, Veterinary Medical Officer; Kelly Helmick, D.V.M.; Kali A. Holder, D.V.M.; David Olsen, Veterinary Technician; and Pattie Walsh, Pathology Technician. We also thank Jennifer Donato, Registrar, Smithsonian’s National Zoo and Conservation Biology Institute (NZCBI), for processing the Biological Materials Transfer Agreement (BMTA) between NZCBI and the University of Louisville Sequencing Technology Center. We are grateful to Brandie Smith, Ph.D., Director of NZCBI, for authorizing the BMTA. Figures were generated in BioRender.

## Declaration of generative AI and AI-assisted technologies in the manuscript preparation process

During the preparation of this work, the author(s) used ChatGPT for improvement of the text. The author(s) reviewed and edited the output as needed and take full responsibility for the content of the published article.

## Methods

### Domestic ferret samples

Ferret experiments were conducted in a BSL2 facility at the University of Pittsburgh in compliance with the guidelines of the Institutional Animal Care and Use Committee (approved protocols 19075697 and 22061230). Ketamine and xylazine were used for sedation for all terminal procedures, followed by cardiac administration of euthanasia solution. Approved University of Pittsburgh Division of Laboratory Animal Resources (DLAR) staff administered euthanasia at time of sacrifice.

Seven adult animals were used for this study. Kidney tissue from a single individual was harvested and flash frozen for long-read genome sequencing. From two animals 131 days post A/Texas/36/1991 (H1N1) influenza infection, blood samples were collected using SST tubes for serum collection. From a single individual, a small blood sample was collected from using a CPT tube for PBMC collection. From the remaining three animals, terminal sampling was performed. Terminal blood samples were collected from using CPT tubes for PBMC collection. Briefly, tubes were inverted gently 5–10 times immediately after drawing and centrifuged at 1,700 × g for 30 minutes at room temperature in a swinging-bucket rotor. The whitish mononuclear cell layer (buffy coat) above the gel plug was collected. The isolated cells were washed twice with PBS, counted, and cryopreserved in a media containing 90% fetal bovine serum and 10% dimethyl sulfoxide (DMSO). 1 cm2 pieces of ferret spleen were collected from three animals at the time of necropsy and placed in 2 mL of R10 buffer (RPMI 1640 medium supplemented with 10% FBS, 100 U/ml penicillin, 100 μg/ml streptomycin, and 2 mM l-glutamine) on ice. The spleen was placed in a 70 um Nylon cell strainer over a 50 mL conical tube and using the rubber tip from a 3 mL syringe, the tissue was completely ground through the strainer. 5 mL of R10 media was used to wash the strainer until no pink tissue remained, at which point the cells were centrifuged at 300 x g for 5 minutes. The RBCs in the cell pellet were lysed with 2 mL of RBC lysis buffer. The remaining splenocytes were washed twice with R10 media, counted, and cryopreserved in a media containing 90% fetal bovine serum and 10% dimethyl sulfoxide (DMSO).

### Samples of other carnivores

Samples of the black-footed ferret (n=1), cheetah (n=1), gray wolf (n=1), maned wolf (n=1), red panda (n=1), and sloth bear (n=1) were obtained from the Biorepository at the Smithsonian’s National Zoo and Conservation Biology Institute (**Table S3**). Samples were originally collected under approved animal care and use protocols as part of routine veterinary procedures, clinical care, or postmortem tissue collection. No live animal experiments or additional animal handling were performed specifically for this study. Sample collection and biobanking procedures were conducted under approved Institutional Animal Care and Use Committee (IACUC) protocols at the originating institutions. All procedures complied with relevant institutional and federal guidelines for work involving endangered species.

### Whole-genome sequencing

High-molecular-weight (HMW) genomic DNA was extracted from samples of the black-footed ferret (lung), domestic ferret (kidney), cheetah (whole blood), red panda (spleen), sloth bear (heart), gray wolf (spleen), and maned wolf (spleen) using multiple protocols optimized per sample (**Table S3**). Primary extraction was performed using the Monarch® HMW DNA Extraction Kit for Tissue or Blood/Cells (New England Biolabs), with alternative extraction using the MagAttract HMW DNA Kit (Qiagen) when necessary to improve yield or quality for specific tissues. In some cases, multiple extraction methods and tissues were evaluated iteratively to obtain sufficient DNA mass and fragment length for long-read sequencing.

DNA quality was assessed using spectrophotometry (NanoVue Plus, Biochrom) for purity (A260/280 and A260/230 ratios), fluorometric quantification (Qubit dsDNA High Sensitivity Assay, Thermo Fisher), and fragment size distribution using the Agilent Femto Pulse system with the Genomic DNA 165 kb kit. To enrich for long DNA fragments, genomic DNA was subjected to size selection using either the Short Read Eliminator (SRE) HT kit (Pacific Biosciences, 103-124-500) to remove fragments <25 kb or BluePippin size selection (Sage Science) with cutoffs ranging from 8–10 kb. In some cases, SRE treatment was omitted due to limited DNA input or sample-specific constraints. Size-selected DNA was sheared to a target fragment size of ∼15–20 kb using g-TUBEs (Covaris) with repeated centrifugation. Sheared DNA was purified using SMRTbell cleanup beads (Pacific Biosciences, 102-158-300) and reassessed for concentration and fragment size. For samples exhibiting non-uniform shearing, additional dilution and re-shearing steps were performed to improve fragment size consistency.

SMRTbell libraries were prepared using the SMRTbell Prep Kit 3.0 (Pacific Biosciences, 102-182-700) following manufacturer protocols with minor modifications to accommodate sample-specific conditions (e.g., input mass, shearing variability). Libraries were quantified using Qubit and assessed on the Femto Pulse. Post-ligation size selection was performed using BluePippin (8–10 kb cutoff) or AMPure PB bead-based selection to remove short fragments and enrich for long inserts. Sequencing libraries were annealed to sequencing primers and bound to polymerase using the Revio Polymerase Kit (Pacific Biosciences, 102-817-600). Libraries were loaded at concentrations ranging from ∼220–390 pM onto 25M SMRT cells (102-202-200) and sequenced on the PacBio Revio platform (102-090-600) using 30-hour movie collections with adaptive loading enabled. Primary processing to generate HiFi reads was performed on-instrument using SMRT Link software, and resulting reads were exported for downstream analysis.

### Genome annotation and inferring orthology

A *de novo* repeat library was created using RepeatModeler v.2.0.7 (flag `--engine NCBI`) on the domestic ferret *Mustela putorius furo* genome assembly. Repeats in the assembly were then soft-masked with RepeatMasker (v.4.2.1; flag `--engine crossmatch -s`). We annotated our assembly using the projection-based annotation pipeline TOGA1 (Tool to Infer Orthologs from Genome Alignments), with default parameters (Kirilenko et al. 2023). Orthologs in the domestic ferret assembly were inferred using 36,664 transcripts and 19,456 coding genes from the GENCODE v.38 (Flicek et al., 2014) annotation of the human genome (hg38) as a reference. Local alignments generated by LASTZ were prepared as TOGA inputs with human-to-other-mammal parameters (K=2400, L=3000, Y=9400, H=2000) and chaining was performed using axtChain with default parameters and a medium gap penalty. Local alignments were improved using RepeatFiller and chainCleaner with default parameters. The completeness of the annotation was assessed using an *a priori* set of genes highly conserved across mammals using compleasm (v.0.2.7; flag `--lineage odb10_mammalia`) in protein mode.

### Generation of the germline domestic ferret IG gene set

IgDetective (v1.1.0) was used to identify contigs containing IG heavy and light chain loci as well to predict putative V, J, and D genes (Sirupurapu et al., 2022). Additionally, variable, joining, and constant genes for heavy, lambda, and kappa loci from all species including those for the domestic ferret within the IMGT^®^ of The International Immunogenetics Information System (Lefranc, 2003) as well as those published (Wong et al., 2020) were mapped onto corresponding contigs using Geneious Prime 2021.1.1 (https://www.geneious.com). Candidate gene segments (V, D, J, and C) were categorized as functional, open reading frame (ORF), and pseudogenes according to “Functionality” of IMGT. Leader sequences were manually annotated. Recombination sequence signals (RSS) were identified using https://www.itb.cnr.it/rss/analyze.html and filtered based on previously described RIC score thresholds (Cowell et al., 2002; Merelli et al., 2010).

To evaluate the concordance between the generated IG germline gene sets and previously published IG genes of the domestic ferret (Lefranc, 2003; Wong et al., 2020; Hebert et al., 2025), V, D, J, and C genes were analyzed across IGH, IGK, and IGL loci. For each detected IG gene, the closest existing IG gene was computed across all four databases. The sequence similarity between two sequences was measured as the percentage of matching nucleotides relative to the length of the shorter sequence. In total, our annotation contained 391 IG genes, of which 350 (89.5%) showed 100% sequence identity to previously published genes. Perfect matches were observed for 81.1% of IGHV genes, 88.4% of IGKV genes, 99.2% of IGLV genes, and 75% of IGLC genes. Also, all IGHD, IGHJ, IGKJ, IGKC, and IGLJ genes matched previously reported sequences exactly (**Figure S8A**). Among genes without perfect matches, 20 IGHV genes showed sequence similarities ranging from 97.92% to 99.66%, six IGHC genes ranged from 99.60% to 99.92% similarity, 13 IGKV genes ranged from 97.04% to 99.66%, one IGLV gene showed 99.66% similarity, and one IGLC gene showed 94.90% similarity to the closest previously reported gene (**Figure S8A**). Previously published genes that were not present in our annotation include 14 IGHV genes (97.62–99.66% similarity to the closest detected gene), one IGHD gene (92.86%), one IGHJ gene (93.75%), one IGHC gene (99.85%), and four IGKV genes (86.01–99.34%) (**Figure S8A**). **Figures S8B** and **S9** visualize the exact overlaps among the four annotations and show that 67.4% of the genes in all annotations were detected at least twice and 54.5% of the genes were detected three or more times. The long-CDRL1 IGLV gene is present in all three analyzed germline IG sets under the names IGLV1-73*01 in Lefranc, 2003; I-LV1_(GL897285.1) in Wong et al., 2020; IGLV1-76*01 in Hebert et al., 2025, as well as in the germline IG gene generated in this study.

### Preparation and sequencing of Iso-Seq libraries

Total RNA was extracted from frozen whole blood or tissue samples using either the AllPrep DNA/RNA Mini Kit (Qiagen, 80204) or the RNeasy Plus Mini Kit (Qiagen, 74134) according to the manufacturer’s instructions. Initial extractions were performed with the AllPrep kit to simultaneously isolate RNA and genomic DNA; however, because genomic DNA quality was suboptimal, subsequent RNA extractions were carried out using the RNeasy Plus kit. RNA purity was assessed using a NanoVue Plus spectrophotometer (Biochrom), and RNA concentration was measured with the Qubit RNA High Sensitivity Assay (Thermo Fisher Scientific, Q32855). RNA integrity was evaluated using either the Agilent 2100 Bioanalyzer with RNA 6000 Nano or Pico kits or the Agilent 4150 TapeStation system with HS RNA ScreenTape. Prior to library preparation, RNA samples selected for sequencing were further purified using RNAclean XP beads (Beckman Coulter, A63987) following the manufacturer’s protocol.

Full-length barcoded cDNA libraries were generated from 300 ng of total RNA using the Iso-Seq Express Template Preparation protocol and the Kinnex Full-Length RNA Kit (Pacific Biosciences, 103-238-700, Rev07 JUN2025) in combination with the Iso-Seq Express 2.0 Kit (Pacific Biosciences, 103-071-500). Amplified cDNA was quantified using the Qubit 1X dsDNA HS Assay Kit (Thermo Fisher Scientific, Q33231) and assessed on the Agilent 4150 TapeStation system with HS D5000 ScreenTape.

For Kinnex library preparation, barcoded cDNA samples were pooled in equimolar amounts, and 55 ng of pooled cDNA was used to prepare concatenated Kinnex libraries using the Kinnex PCR 8-Fold Kit (Pacific Biosciences, 103-071-600) and the Kinnex Concatenation Kit (Pacific Biosciences, 103-071-800) according to the manufacturer’s protocol. Final libraries were quantified with Qubit and evaluated on the Agilent 4150 TapeStation system using HS Genomic DNA ScreenTape.

Sequencing libraries were prepared at a loading concentration of 225 pM using calculations generated in SMRT Link v25.2 (Pacific Biosciences). Libraries were annealed to the Kinnex sequencing primer and bound to polymerase using the Revio SPRQ Polymerase Kit (Pacific Biosciences, 103-496-900), followed by purification with SMRTbell cleanup beads (Pacific Biosciences, 102-158-300). Sequencing was performed on a single 25M SMRT Cell (102-202-200) on the PacBio Revio platform (102-090-600) with a 24-hour movie collection time and adaptive loading enabled. Primary analysis and generation of HiFi reads were performed on the instrument using SMRT Link software before export to a local server for downstream analyses. Sequencing was carried out at the University of Louisville Sequencing Technology Center (Kentucky, USA).

### Processing raw Iso-Seq reads

Raw subread BAM files were processed using the PacBio Iso-Seq pipeline. First, BAM indices were generated using `pbindex`. Reads were then segmented using `skera` split with MAS-Seq adapter sequences provided in `mas8_primers.fasta`. Primer trimming and demultiplexing were performed using the lima tool with the `--isoseq`, `--peek-guess`, and `--split-named` options and the PacBio Iso-Seq v2 primer set (`IsoSeq_v2_primers_12.fasta`). Finally, full-length non-concatemer reads were refined for each barcode separately using `isoseq refine --require-polya` to retain polyadenylated transcripts. The files `mas8_primers.fasta` and `IsoSeq_v2_primers_12.fasta` were obtained from the PacBio documentation and support resources. The clustering stage (`isoseq cluster`) was intentionally omitted to avoid collapsing highly similar V(D)J transcripts.

### Preparation and sequencing of Rep-Seq libraries for domestic ferrets

RNA was isolated from ferret peripheral blood mononuclear cells (PBMCs) and splenocytes using the Zymogen Quick-DNA/RNA Miniprep Kit (Zymogen, cat. no. D7001) and 5’RACE cDNA was generated using the SMARTer 3’/5’RACE Kit (Takara, cat. no. 634859). Published nested IGLC gene primers (Wong et al. 2020) were used for amplification of IgL sequences. The first round PCR was conducted with the SeqAmp polymerase (Takara Cat #638504) and associated buffer and universal primer mix from the SMARTer Kit, 2.5 µL of 5’RACE cDNA, 10 µM outer IgL primer, with an initial denaturation of 94°C for 1 minute, then 40 cycles of 94°C for 30s, 68°C for 45s, and 72°C for 3 minutes and final extension of 5 minutes. The PCR product from round 1 was diluted 1:30 in TE buffer and 5uL used as template for a second round of a three-step PCR with 20 cycles of 94°C for 30s, 68°C for 45s, and 72°C for 3 minutes and final extension of 5 minutes. A dual sided SPRIselect (Beckman Coulter, cat. no. B23317) bead size selection was performed on PCR products. Libraries were barcoded using the NEBNext Ultra II DNA Library Prep Kit for Illumina^®^ (New England Biolabs, #E7645S and #E7370L). Libraries were pooled and sequenced using a MiSeq Reagent Kit v3 600 cycle (Illumina, cat. no. MS-102-3003) on an Illumina MiSeq instrument.

### Preparation and sequencing of scRNA-Seq libraries for domestic ferrets

PBMCs were thawed, washed once with Fetal Bovine Solution, counted, and resuspended in a 1x phosphate-buffered saline + 0.2% bovine serum albumin. Approximately 20,000 cells were loaded per 10x lane. The Chromium GEM Single cell 5’ kit v2 (10x Genomics PN-1000374) was used and protocol Rev D was followed for the preparation of cDNA and gene expression library generation. For single cell BCR library generation, published IG primers were used (Walsh et al., 2025). Briefly, first round BCR amplification gene specific primers were used at 0.5 µM final concentration with 1 µM final concentration of 10x compatible forward primer in TE buffer and 12 cycles of amplification were used. Second round BCR amplification gene specific primers were used at 0.5 µM final concentration with 1 µM final concentration of 10x compatible forward primer in the TE buffer and 10 cycles of amplification were used. After library prep, quality control was performed using an Agilent 2100 Bioanalyzer (Agilent Technologies, cat. no. G2939B) and quantification, pooling and dilution of libraries to 1.5 nM was performed using the KAPA Library Quantification Kit (Roche, cat. no. 07960140001). Libraries were pooled and sequenced using a Nova-seq SP 200 cycle kit (Illumina, San Diego, CA; cat. no. 20040719) on an Illumina NovaSeq 6000 instrument.

### Monoclonal production

Recombinant monoclonal antibody IgGs were expressed in Expi293F cells via a 1:1 co-transfection of plasmids encoding the respective heavy and light chains, or a 1:1:1 co-transfection including a pcDNA3.1 plasmid encoding human tyrosylprotein sulfotransferase 2 (TPST2; UniProt O60704). Transfection was performed using the ExpiFectamine transfection kit (Thermo Fisher Scientific, cat. no. A14525) following the manufacturer’s protocols. In brief, cells were cultured and maintained at 37°C, 8% CO2. On the day of transfection, cells were diluted to 2.5 x 10^6 cells per mL and transfected with DNA at a 1 µg per mL ratio. ExpiFectamine transfection enhancers were added 20 hours post transfection, and cell supernatants were harvested and filtered 4 days post transfection. Filtered supernatants were subsequently run over HiTrap MabSelect SuRe columns (Cytiva) to capture expressed IgGs. Eluted IgGs were concentrated using Amicon ultra-15 centrifugal filters with a 30 kDa molecular weight cut off (MWCO) (Millipore, cat. no. UFC9030) and purified via size exclusion chromatography (SEC) using a Superdex 200 Increase 10/300 GL column (Cytiva) into 1x Tris-buffered saline supplemented with sodium azide (TBS-Az: 20 mM Tris-HCl pH 8.0, 150 mM sodium chloride, 0.02% (v/v) sodium azide).

### Western blot

Western blot analysis was performed to assess protein sulfation. Samples included 1 µg of a control sulfated Fab, 0.5 µg of a control, non-sulfated monoclonal antibody, and 0.5 µg of a ferret monoclonal antibody (both non-reduced and reduced forms). For reduction, ferret monoclonal antibody samples were treated with 20 µM dithiothreitol (DTT) prior to loading. Proteins were separated on a 12% polyacrylamide gel (Thermo Fisher Scientific NP0329BOX) under denaturing conditions and dry transferred to a polyvinylidene difluoride (PVDF) membrane (Thermo Fisher Scientific IB24002). The membrane was blocked for 1 hour at room temperature in 5% bovine serum albumin (BSA) dissolved in phosphate-buffered saline with 0.1% Tween-20 (PBST). Following blocking, the membrane was incubated overnight at room temperature with an anti-sulfotyrosine antibody (Abcam, ab136481) diluted 1:500 in 5% BSA + PBST. The membrane was then washed five times with PBST, followed by incubation with an anti-mouse secondary antibody (LI-COR Biosciences #925-68070) at 1:10,000 at room temperature for 1 hour. After secondary antibody incubation, the membrane was washed five additional times with PBST. Images were captured using a Bio-Rad ChemiDoc Touch Imaging System (Bio-Rad Laboratories, cat. no. 1708370).

### Biolayer interferometry

Sulfation was assessed using biolayer interferometry (BLI) on an Octet RED96 system (ForteBio/Sartorius). All experiments were conducted at 25°C. Octet AHC biosensor (Sartorius, #18-5060) were hydrated in PBS w/ 0.1 % Tween-20 for at least 20 minutes prior to use. Humanized ferret monoclonals were immobilized onto Streptavidin biosensor tips at a concentration of 10 µg/mL for 2 minutes. Monoclonal loading was followed by a brief wash in an assay buffer to remove unbound material. Sulfation was evaluated by immersing the monoclonal-loaded tips into wells containing - anti-sulfotyrosine antibody (Sulfo-1C-A2, EMD Millipore #05-1100X) at 1:100 dilution in phosphate-buffered saline w/ 0.1 % Tween 20. Association phase was recorded for 2 minutes and dissociation was recorded for 4 minutes each. Between monoclonals, biosensor tips were regenerated using 10 mM glycine (pH 2.0) for 3 cycles. A control sample containing wells containing buffer alone and a non-sulfated human monoclonal were included to subtract non-specific binding and baseline drift.

### Macrophage infections

RAW 264.7 cells were seeded into 24-well tissue culture plates at 2×10^5^ and incubated at 37°C, 5% CO_2_. *Y. pestis* KIM1001 (pCD1+pgm-) (Palace et al., 2018) was cultured in BHI broth overnight at 26°C, then sub-cultured 1:10 at 26°C and 37°C for 3 hours prior to infections. *Y. pestis* (2×10^7^ colony forming units (CFU)/mL) was incubated with either 5% serum or 100 µg/mL monoclonal antibody for 20 minutes at 37°C with rotation. RAW 264.7 cells were infected with a multiplicity of infection (MOI) of 10 with either untreated *Y. pestis* alone (26°C and 37°C), or *Y. pestis* (37°C) pre-incubated with serum or monoclonal antibody and incubated at 37°C, 5% CO_2_. After 20 minutes, extracellular bacteria were killed by adding 16 µg/ml gentamicin to DMEM media. After 1 hour, the media was replaced with DMEM containing 10% FBS and 4 ug/mL gentamicin. At 2 and 6 hours post-infection, cells were washed twice with 1X PBS and lysed with 1% Triton X-100. Lysates were serially diluted and plated in triplicate on BHI agar to quantify CFU. Phagocytosis was defined as the percentage of CFU recovered from cell lysates 2 hours post-infection relative to the initial inoculum. Intracellular survival was defined as the percentage of CFU recovered at 6 hours relative to 2 hours post-infection.

## References

Abayev M, Rodrigues JP, Srivastava G, Arshava B, Jaremko Ł, Jaremko M, Naider F, Levitt M, Anglister J. The solution structure of monomeric CCL 5 in complex with a doubly sulfated N-terminal segment of CCR 5. The FEBS journal. 2018 Jun;285(11):1988–2003.

Abbott RC, Osorio JE, Bunck CM, Rocke TE. Sylvatic plague vaccine: a new tool for conservation of threatened and endangered species?. EcoHealth. 2012 Sep;9(3):243–50.

Acharya P, Dogo-Isonagie C, LaLonde JM, Lam SN, Leslie GJ, Louder MK, Frye LL, Debnath AK, Greenwood JR, Luongo TS, Martin L. Structure-based identification and neutralization mechanism of tyrosine sulfate mimetics that inhibit HIV-1 entry. ACS chemical biology. 2011 Aug 5;6(10).

Achtman M, Zurth K, Morelli G, Torrea G, Guiyoule A, Carniel E. Yersinia pestis, the cause of plague, is a recently emerged clone of Yersinia pseudotuberculosis. Proceedings of the National Academy of Sciences. 1999 Nov 23;96(24):14043–8.

Alonzo III F, Kozhaya L, Rawlings SA, Reyes-Robles T, DuMont AL, Myszka DG, Landau NR, Unutmaz D, Torres VJ. CCR5 is a receptor for Staphylococcus aureus leukotoxin ED. Nature. 2013 Jan 3;493(7430):51-5.

Alt FW, Baltimore D. Joining of immunoglobulin heavy chain gene segments: implications from a chromosome with evidence of three D-JH fusions. Proceedings of the National Academy of Sciences. 1982 Jul;79(13):4118–22.

Avnir Y, Watson CT, Glanville J, Peterson EC, Tallarico AS, Bennett AS, Qin K, Fu Y, Huang CY, Beigel JH, Breden F. IGHV1-69 polymorphism modulates anti-influenza antibody repertoires, correlates with IGHV utilization shifts and varies by ethnicity. Scientific reports. 2016 Feb 16;6(1):20842.

Barbieri R, Signoli M, Chevé D, Costedoat C, Tzortzis S, Aboudharam G, Raoult D, Drancourt M. Yersinia pestis: the natural history of plague. Clinical microbiology reviews. 2020 Dec 16;34(1):10–128.

Bernard O, Hozumi N, Tonegawa S. Sequences of mouse immunoglobulin light chain genes before and after somatic changes. Cell. 1978 Dec 1;15(4):1133–44.

Bevins SN, Chandler JC, Barrett N, Schmit BS, Wiscomb GW, Shriner SA. Plague exposure in mammalian wildlife across the Western United States. Vector-Borne and Zoonotic Diseases. 2021 Sep;21(9):667–74.

Biggins DE, Kosoy MY. Influences of introduced plague on North American mammals: implications from ecology of plague in Asia. Journal of Mammalogy. 2001 Nov 1;82(4):906–16.

Bolotin DA, Poslavsky S, Mitrophanov I, Shugay M, Mamedov IZ, Putintseva EV, Chudakov DM. MiXCR: software for comprehensive adaptive immunity profiling. Nature methods. 2015 May;12(5):380–1.

Burns R, Williams ES, O’Toole D, Dubey JP. Toxoplasma gondii infections in captive black-footed ferrets (Mustela nigripes), 1992–1998: clinical signs, serology, pathology, and prevention. The Journal of Wildlife Diseases. 2003 Oct 1;39(4):787–97.

Cheng H, Concepcion GT, Feng X, Zhang H, Li H. Haplotype-resolved de novo assembly using phased assembly graphs with hifiasm. Nature methods. 2021 Feb;18(2):170–5.

Choe H, Li W, Wright PL, Vasilieva N, Venturi M, Huang CC, Grundner C, Dorfman T, Zwick MB, Wang L, Rosenberg ES. Tyrosine sulfation of human antibodies contributes to recognition of the CCR5 binding region of HIV-1 gp120. Cell. 2003 Jul 25;114(2):161–70.

Cowell LG, Davila M, Kepler TB, Kelsoe G. Identification and utilization of arbitrary correlations in models of recombination signal sequences. Genome biology. 2002 Nov 21;3(12):research0072-1.

Crooks GE, Hon G, Chandonia JM, Brenner SE. WebLogo: a sequence logo generator. Genome research. 2004 Jun 1;14(6):1188–90.

Cullen, J.N., Martin, J., Vilella, A.J., Treeful, A., Sargan, D., Bradley, A. and Friedenberg, S.G., 2022. Development and application of a next-generation sequencing protocol and bioinformatics pipeline for the comprehensive analysis of the canine immunoglobulin repertoire. Plos one, 17(7), p.e0270710.

Cunningham MW. Molecular mimicry. In: Encyclopedia of Life Sciences. Chichester (UK): John Wiley & Sons, Ltd; 2009.

Dorfman T, Moore MJ, Guth AC, Choe H, Farzan M. A tyrosine-sulfated peptide derived from the heavy-chain CDR3 region of an HIV-1-neutralizing antibody binds gp120 and inhibits HIV-1 infection. Journal of Biological Chemistry. 2006 Sep 29;281(39):28529–35.

Ducancel F, Mérienne K, Fromen-Romano C, Trémeau O, Pillet L, Drevet P, Zinn-Justin S, Boulain JC, Ménez A. Mimicry between receptors and antibodies: Identification of snake toxin determinants recognized by the acetylcholine receptor and an acetylcholine receptor-mimicking monoclonal antibody. Journal of Biological Chemistry. 1996 Dec 6;271(49):31345–53.

Engelbrecht E, Rodriguez OL, Lees W, Vanwinkle Z, Shields K, Schultze S, Gibson WS, Smith DR, Jana U, Saha S, Peres A. Germline polymorphisms in the immunoglobulin kappa and lambda loci underpinning antibody light chain repertoire variability. Nature Communications. 2025 Nov 28.

Farzan M, Mirzabekov T, Kolchinsky P, Wyatt R, Cayabyab M, Gerard NP, Gerard C, Sodroski J, Choe H. Tyrosine sulfation of the amino terminus of CCR5 facilitates HIV-1 entry. Cell. 1999 Mar 5;96(5):667–76.

Fields BA, Goldbaum FA, Ysern X, Poijak RJ, Mariuzza RA. Molecular basis of antigen mimicry by an anti-idiotope. Nature. 1995 Apr 20;374(6524):739–42.

Flicek P, Amode MR, Barrell D, Beal K, Billis K, Brent S, Carvalho-Silva D, Clapham P, Coates G, Fitzgerald S, Gil L. Ensembl 2014. Nucleic acids research. 2014 Jan 1;42(D1):D749–55.

Gage KL, Kosoy MY. Natural history of plague: perspectives from more than a century of research. Annu Rev Entomol. 2005;50:505–28.

Godbey JL, Biggins DE, Garelle D. Exposure of captive black-footed ferrets to plague and implications for species recovery. Recovery of the black-footed ferret: progress and continuing challenges. US Geological Survey Scientific Investigations Report. Fort Collins, Colorado. 2006:233–7.

Hebert LS, Pickens W, Satterwhite E, Soto GB, Pflaum FM, Zhan M, Moody MA, Kain J, Kirchenbaum GA, Ferguson JA, Langel SN. Gene discovery and expression analysis of the B cell receptor repertoire in the domestic ferret model. Vaccine. 2025 Oct 3;64:127725.

Heinrich SK, Wachter B, Aschenborn OH, Thalwitzer S, Melzheimer J, Hofer H, Czirják GÁ. Feliform carnivores have a distinguished constitutive innate immune response. Biology open. 2016 May 15;5(5):550–5.

Henikoff S, Henikoff JG. Amino acid substitution matrices from protein blocks. Proceedings of the national academy of sciences. 1992 Nov 15;89(22):10915–9.

Hozumi N, Tonegawa S. Evidence for somatic rearrangement of immunoglobulin genes coding for variable and constant regions. Proceedings of the National Academy of Sciences. 1976 Oct;73(10):3628–32.

Huang, C.C., Venturi, M., Majeed, S., Moore, M.J., Phogat, S., Zhang, M.Y., Dimitrov, D.S., Hendrickson, W.A., Robinson, J., Sodroski, J. and Wyatt, R. Structural basis of tyrosine sulfation and VH-gene usage in antibodies that recognize the HIV type 1 coreceptor-binding site on gp120. Proceedings of the National Academy of Sciences. 2004. 101(9), pp.2706–2711.

Huang CC, Lam SN, Acharya P, Tang M, Xiang SH, Hussan SS, Stanfield RL, Robinson J, Sodroski J, Wilson IA, Wyatt R. Structures of the CCR5 N terminus and of a tyrosine-sulfated antibody with HIV-1 gp120 and CD4. Science. 2007 Sep 28;317(5846):1930–4.

Ibrahim HM, Xuan X, Nishikawa Y. Toxoplasma gondii cyclophilin 18 regulates the proliferation and migration of murine macrophages and spleen cells. Clinical and vaccine immunology. 2010 Sep;17(9):1322–9.

Ivanov MI, Hill J, Bliska JB. Direct neutralization of type III effector translocation by the variable region of a monoclonal antibody to Yersinia pestis LcrV. Clin Vaccine Immunol. 2014 May;21(5):667–73.

Jachowski D. Wild Again: The Struggle to Save the Black-Footed Ferret. 2014. University of California Press.

Jen CH, Moore KL, Leary JA. Pattern and temporal sequence of sulfation of CCR5 N-terminal peptides by tyrosylprotein sulfotransferase-2: an assessment of the effects of N-terminal residues. Biochemistry. 2009 Jun 16;48(23):5332–8.

Johnson TA, Mashimo Y, Wu JY, Yoon D, Hata A, Kubo M, Takahashi A, Tsunoda T, Ozaki K, Tanaka T, Ito K. Association of an IGHV3-66 gene variant with Kawasaki disease. Journal of human genetics. 2021 May;66(5):475–89.

Kabat EA, Wu TT. Attempts to locate complementarity-determining residues in the variable positions of light and heavy chains. Annals of the New York Academy of Sciences. 1971 Dec;190(1):382–93.

Kirilenko BM, Munegowda C, Osipova E, Jebb D, Sharma V, Blumer M, Morales AE, Ahmed AW, Kontopoulos DG, Hilgers L, Lindblad-Toh K. Integrating gene annotation with orthology inference at scale. Science. 2023 Apr 28;380(6643):eabn3107.

Lee PS, Ohshima N, Stanfield RL, Yu W, Iba Y, Okuno Y, Kurosawa Y, Wilson IA. Receptor mimicry by antibody F045–092 facilitates universal binding to the H3 subtype of influenza virus. Nature communications. 2014 Apr 10;5(1):3614.

Lefranc MP. IMGT, the international ImMunoGeneTics database®. Nucleic acids research. 2003 Jan 1;31(1):307–10.

Li H. Minimap2: pairwise alignment for nucleotide sequences. Bioinformatics. 2018 Sep 15;34(18):3094–100.

Mahmoudi A, Kryštufek B, Sludsky A, Schmid BV, De Almeida AM, Lei X, Ramasindrazana B, Bertherat E, Yeszhanov A, Stenseth NC, Mostafavi E. Plague reservoir species throughout the world. Integrative Zoology. 2021 Nov;16(6):820–33.

Matchett MR, Biggins DE, Carlson V, Powell B, Rocke T. Enzootic plague reduces black-footed ferret (Mustela nigripes) survival in Montana. Vector-Borne and Zoonotic Diseases. 2010 Feb;10(1):27–35.

Mecsas J, Franklin G, Kuziel WA, Brubaker RR, Falkow S, Mosier DE. CCR5 mutation and plague protection. Nature. 2004 Feb 12;427(6975):606.

Merelli I, Guffanti A, Fabbri M, Cocito A, Furia L, Grazini U, Bonnal RJ, Milanesi L, McBlane F. RSSsite: a reference database and prediction tool for the identification of cryptic Recombination Signal Sequences in human and murine genomes. Nucleic acids research. 2010 Jul 1;38(suppl_2):W262–7.

Miller MA, Moriarty ME, Henkel L, Tinker MT, Burgess TL, Batac FI, Dodd E, Young C, Harris MD, Jessup DA, Ames J. Predators, disease, and environmental change in the nearshore ecosystem: mortality in southern sea otters (Enhydra lutris nereis) from 1998–2012. Frontiers in Marine Science. 2020 Nov 19;7:582.

Mize EL, Grassel SM, Britten HB. Fleas of black-footed ferrets (Mustela nigripes) and their potential role in the movement of plague. J Wildl Dis. 2017 Jul;53(3):521–531.

Monigatti F, Gasteiger E, Bairoch A, Jung E. The Sulfinator: predicting tyrosine sulfation sites in protein sequences. Bioinformatics. 2002 May;18(5):769–70.

Moore RM, Harrison AO, McAllister SM, Polson SW, Wommack KE. Iroki: automatic customization and visualization of phylogenetic trees. PeerJ. 2020 Feb 26;8:e8584.

Osei-Owusu P, Charlton TM, Kim HK, Missiakas D, Schneewind O. FPR1 is the plague receptor on host immune cells. Nature. 2019 Oct 3;574(7776):57–62.

Ota T, Nei M. Divergent evolution and evolution by the birth-and-death process in the immunoglobulin VH gene family. Molecular biology and evolution. 1994 May 1;11(3):469–82.

Oyston PC, Williamson D. Plague: infections of companion animals and opportunities for intervention. Animals. 2011 Jun 21;1(2):242–55.

Palace SG, Proulx MK, Szabady RL, Goguen JD. Gain-of-Function Analysis Reveals Important Virulence Roles for the Yersinia pestis Type III Secretion System Effectors YopJ, YopT, and YpkA. Infect Immun. 2018 Aug 22;86(9):e00318–18.

Parks T, Mirabel MM, Kado J, Auckland K, Nowak J, Rautanen A, Mentzer AJ, Marijon E, Jouven X, Perman ML, Cua T. Association between a common immunoglobulin heavy chain allele and rheumatic heart disease risk in Oceania. Nature communications. 2017 May 11;8(1):14946.

Pennell M, Rodriguez OL, Watson CT, Greiff V. The evolutionary and functional significance of germline immunoglobulin gene variation. Trends in immunology. 2023 Jan 1;44(1):7–21.

Peres A, Upadhyay AA, Klein V, Saha S, Rodriguez OL, Vanwinkle ZM, Karunakaran K, Metz A, Lauer W, Lin MC, Melton T. Population-level genomic analysis of immunoglobulin loci variation in rhesus macaques reveals extensive germline diversity. Immunity. 2026 Jan 13;59(1):213–28.

Pospelova M, Voss K, Zamyatin A, Watson CT, Koepfli KP, Bankevich A, Pennell M, Safonova Y. Comparative analysis of mammalian adaptive immune loci revealed spectacular divergence and common genetic patterns. Molecular biology and evolution. 2025 Jul;42(7):msaf152.

Rasmussen S, Allentoft ME, Nielsen K, Orlando L, Sikora M, Sjögren KG, Pedersen AG, Schubert M, Van Dam A, Kapel CM, Nielsen HB. Early divergent strains of Yersinia pestis in Eurasia 5,000 years ago. Cell. 2015 Oct 22;163(3):571–82.

Rascovan N, Sjögren KG, Kristiansen K, Nielsen R, Willerslev E, Desnues C, Rasmussen S. Emergence and spread of basal lineages of Yersinia pestis during the Neolithic decline. Cell. 2019 Jan 10;176(1):295–305.

Rodriguez OL, Safonova Y, Silver CA, Shields K, Gibson WS, Kos JT, Tieri D, Ke H, Jackson KJ, Boyd SD, Smith ML. Genetic variation in the immunoglobulin heavy chain locus shapes the human antibody repertoire. Nature communications. 2023 Jul 21;14(1):4419.

Salkeld DJ, Stapp P. Seroprevalence rates and transmission of plague (Yersinia pestis) in mammalian carnivores. Vector-Borne and Zoonotic Diseases. 2006 Sep;6(3):231–9.

Seibert C, Cadene M, Sanfiz A, Chait BT, Sakmar TP. Tyrosine sulfation of CCR5 N-terminal peptide by tyrosylprotein sulfotransferases 1 and 2 follows a discrete pattern and temporal sequence. Proceedings of the National Academy of Sciences. 2002 Aug 20;99(17):11031–6.

Sheahan KL, Isberg RR. Identification of mammalian proteins that collaborate with type III secretion system function: involvement of a chemokine receptor in supporting translocon activity. MBio. 2015 Feb 27;6(1):10–128.

Shlemov A, Bankevich S, Bzikadze A, Turchaninova MA, Safonova Y, Pevzner PA. Reconstructing antibody repertoires from error-prone immunosequencing reads. The Journal of Immunology. 2017 Nov;199(9):3369–80.

Schmidt AG, Therkelsen MD, Stewart S, Kepler TB, Liao HX, Moody MA, Haynes BF, Harrison SC. Viral receptor-binding site antibodies with diverse germline origins. Cell. 2015 May 21;161(5):1026–34.

Sievers F, Wilm A, Dineen D, Gibson TJ, Karplus K, Li W, Lopez R, McWilliam H, Remmert M, Söding J, Thompson JD. Fast, scalable generation of high-quality protein multiple sequence alignments using Clustal Omega. Molecular systems biology. 2011 Oct 11;7:539.

Sirupurapu V, Safonova Y, Pevzner PA. Gene prediction in immunoglobulin loci. Genome Research. 2022. 32:1152–1169.

Sun YC, Jarrett CO, Bosio CF, Hinnebusch BJ. Retracing the evolutionary path that led to flea-borne transmission of Yersinia pestis. Cell host & microbe. 2014 May 14;15(5):578–86.

Tollenaere C, Rahalison L, Ranjalahy M, Rahelinirina S, Duplantier JM, Brouat C. CCR5 polymorphism and plague resistance in natural populations of the black rat in Madagascar. Infection, Genetics and Evolution. 2008 Dec 1;8(6):891–7.

Totikov AA, Tomarovsky AA, Perelman PL, Bulyonkova TM, Serdyukova NA, Yakupova AR, Mohr D, Foerster DW, Grau Jipoulou JH, Beklemisheva VR, Sidorov M. Comparative genomics and phylogenomics of the Mustelinae lineage (Mustelidae, Carnivora). Genome Biology and Evolution, 2026. 18(3): evag014.

Upham NS, Esselstyn JA, Jetz W. Inferring the mammal tree: species-level sets of phylogenies for questions in ecology, evolution, and conservation. PLoS biology. 2019 Dec 4;17(12):e3000494.

Walker LM, Phogat SK, Chan-Hui PY, Wagner D, Phung P, Goss JL, Wrin T, Simek MD, Fling S, Mitcham JL, Lehrman JK. Broad and potent neutralizing antibodies from an African donor reveal a new HIV-1 vaccine target. Science. 2009 Oct 9;326(5950):285–9.

Walsh ES, Yang K, Tollison TS, Seenu S, Adams N, Zeitoun G, Sideri I, Folch G, Brochu HN, Chou H, Kossida S. Development of ferret immune repertoire reference resources and single-cell-based high-throughput profiling assays. Journal of Virology. 2025 Apr 15;99(4):e00181–25.

Williams ES, Thome ET, Quan TJ, Anderson SL. Experimental infection of domestic ferrets (Mustela putorius furo) and Siberian polecats (Mustela eversmanni) with Yersinia pestis. The Journal of Wildlife Diseases. 1991 Jul 1;27(3):441–5.

Wisely SM, Buskirk SW, Fleming MA, McDonald DB, Ostrander EA. Genetic diversity and fitness in black-footed ferrets before and during a bottleneck. Journal of Heredity. 2002 Jul 1;93(4):231–7.

Wong J, Tai CM, Hurt AC, Tan HX, Kent SJ, Wheatley AK. Sequencing B cell receptors from ferrets (Mustela putorius furo). PLoS One. 2020 May 29;15(5):e0233794.

Ye J, Ma N, Madden TL, Ostell JM. IgBLAST: an immunoglobulin variable domain sequence analysis tool. Nucleic acids research. 2013 Jul 1;41(W1):W34–40.

Yoo D, Rhie A, Hebbar P, Antonacci F, Logsdon GA, Solar SJ, Antipov D, Pickett BD, Safonova Y, Montinaro F, Luo, et al. Complete sequencing of ape genomes. Nature. 2025 May 8;641(8062):401–18.

