## Supplemental Materials for "Disruption of a CCR5-like immunoglobulin gene is linked to plague susceptibility in black-footed ferrets"

1 Computer Science and Engineering Department, Pennsylvania State University, State College, PA 16803, USA.
2 Huck Institutes of the Life Sciences, Pennsylvania State University, State College, PA 16803, USA.
3 Department of Pathology, Stanford University, Stanford, CA 94305, USA.
4 Department of Microbiology & Immunology, Stanford University, Stanford, CA 94305, USA.
5 Department of Microbiology and Immunology, University of Louisville School of Medicine, Louisville, KY 40202, USA.
6 Constructor University Bremen gGmbH, Bremen, 28759, Germany.
7 Department of Biology, Stanford University, Stanford, CA 94305, USA.
8 Department of Computational Biology, Cornell University, Ithaca, NY 14853, USA.
9 Institute for Systems Genetics, NYU Langone Health, New York, NY 10016, USA.
10 Department of Biochemistry and Molecular Pharmacology, NYU Langone Health, New York, NY 10016, USA.
11 Sequencing Technology Center, University of Louisville, Louisville, KY 40292, USA.
12 Department of Biological Sciences, Smith College, Northampton, MA 01063, USA.
13 Department of Biochemistry and Molecular Genetics, University of Louisville, Louisville, KY 40292, USA.
14 Department of Microbiology and Molecular Genetics, University of Pittsburgh School of Medicine, Pittsburgh, PA 15219, USA.
15 Center for Vaccine Research, University of Pittsburgh School of Medicine, Pittsburgh, PA 15261, USA.
16 Department of Microbiology, University of Louisville, Louisville, KY 40292, USA.
17 Freeman Hrabowski Scholar, Howard Hughes Medical Institute, Chevy Chase, MD 20815, USA.
18 Department of Microbiology and Immunology, Emory School of Medicine, Atlanta, GA 30322, USA.
19 Sean N. Parker Center for Allergy and Asthma Research, Stanford University, Stanford, CA 94305, USA.
20 Center for Predictive Medicine for Biodefense and Emerging Infectious Diseases, University of Louisville, Louisville, KY 40202, USA.
21 College of Science, George Mason University, Fairfax, VA 22030, USA.
22 Colossal Biosciences, Austin, TX 78701, USA.
† These authors contributed equally to this work.

### Table of contents

#### Supplemental Figures


- Figure S1. The ferret long-CDRL1 IGLV gene.
- Figure S2. Read depth of the steppe polecat subject from Inner Mongolia, China (ERR7198277) aligned to the genome of the steppe polecat subject from Altai, Russia (GCA_963422785.1).
- Figure S3. Alignment of the long-CDRL1s IGLV genes found in the domestic ferret (top line) and the Northern sea otter (bottom line).
- Figure S4. The distribution of similarity scores between CDRL1s and CCR5s across six Carnivora families: Mustelidae, Phocidae, Ursidae, Canidae, Herpestidae, Felidae.
- Figure S5. Counts of somatic hypermutations (SHMs) in the V gene (A) and CDRL1 (B) of the long-CDRL1 IGL sequences across five carnivoran species: domestic ferret (PBMC and spleen), sloth bear (spleen), domestic dog (PBMC), gray wolf (spleen), and maned wolf (spleen).
- Figure S6. dN/dS values computed based on somatic hypermutations in the CDRL1 (A) and non-CDRL1 parts (B) of the long-CDRL1 IGL sequences across five carnivoran species: domestic ferret (PBMC and spleen), sloth bear (spleen), domestic dog (PBMC), gray wolf (spleen), and maned wolf (spleen).
- Figure S7. 5-mers DSYGY (highlighted in green), DYLDV (highlighted in purple), and YAMGY (yellow) with significantly higher fractions in long-CDRL1 IGL Abs have perfect matches to one IGHD gene and two IGHJ genes of domestic ferret.
- Figure S8. Comparison of domestic ferret IG genes with existing IG annotations.
- Figure S9. Venn diagrams showing IGHV, IGKV, IGLV, IGHC, and IGLC genes shared among four analyzed annotations of domestic ferrets.


#### Supplemental Tables


- Table S1. Population data of Mustelinae species.
- Table S2. Publicly available genomes of carnivoran species selected for the comparative analysis.
- Table S3. Information about samples of six newly sequenced carnivoran species: black-footed ferret, cheetah, gray wolf, maned wolf, red panda, and sloth bear.
- Table S4. Characteristics of the new genome assemblies.
- Table S5. Characteristics of the new domestic ferret Rep-seq IGL libraries.
- Table S6. Characteristics of the new Iso-Seq libraries for the black-footed ferret, red panda, sloth bear, gray wolf, and maned wolf.
- Table S7. Characteristics of publicly available Rep-Seq datasets of the domestic ferrets and domestic dogs used in the study.
- Table S8. Characteristics of publicly available scRNA-Seq data of the domestic ferrets used in the study.

#### Supplemental Figures

**A**

```
H. sapiens (IGLV1-40)      CAGTCTGTGCTGACGCAGCCGCCCTCAGTGTCTGGGGCCCCAGGGCAGAGGGTCACCATC    60
M. putorius furo           TGGGCTGTGCTGACTCAGCCTCCCTATGTGTCTGGGGCCCTAGGTGAGAGTGTCACCATC    54
M. nigripes                TGGGCTGTGCTGACTCAGCCTCTCTCTGTGTCTGGGACCCTA--GGTGAGTGTCACCATC    58
                             * ********** ***** * **  ********* *** *     *** *********

H. sapiens (IGLV1-40)      TCCTGCACTGGGAGCAGCTCCAACATCGGGGC---------------AGGTTATGATGTA    105
M. putorius furo           TCCTGCACTGGAATCCCCACCAGCATAGATTATGATGAAGAGGAATACACATATAATGTG    114
M. nigripes                TCCTGCACTGGAATCCCCACCAGCATAGATTATGATGAAGAGGAATACACATATGATGTG    118
                           *********** * *  * *** *** *                       *** ****

H. sapiens (IGLV1-40)      CACTGGTACCAGCAGCTTCCAGGAACAGCCCCCAAACTCCTCATCTATGGTAACAGCAAT    165
M. putorius furo           AACTGGTACCAACAGCTCCAAGGAAAGGTACCCATTCTACTCATCTATGGAGATAATAAC    174
M. nigripes                AACTGGTACCAACAGCTCCAAGGAAAGGTACCCATTCTACTCATCTATGGAGATAATAAC    178
                            ********** ***** * *****  *  ****  ** ***********  * *  **

H. sapiens (IGLV1-40)      CGGCCCTCAGGGGTCCCTGACCGATTCTCTGGCTCCAAGTCTGGCACCTCAGCCTCCCTG    225
M. putorius furo           AGAAATCCTGGAGTCCCTGATCGATTCTCTGGTTCCAAGTCAGGCAGCTCAGCCTCCCTG    234
M. nigripes                AGAAATCCTGGAGTCCCTGATCGATTCTCTGGTTCCAAGTCAGGCAGCTCAGCCTCCCTG    238
                            *     * ** ******** *********** ******** **** *************

H. sapiens (IGLV1-40)      GCCATCACTGGGCTCCAGGCTGAGGATGAGGCTGATTATTACTGCCAGTCCTATGACAGC    285
M. putorius furo           ACCATCAGTGGCCTTCAGGCTGAAGATGAGGCTGATTATTACTGCCAGTCCACTGACAGT    294
M. nigripes                ACCATCAGTGGCCTTCAGGCTGAAGATGAGGCTGATTATTACTGCCAGTCCACTGACAGT    298
                            ****** *** ** ******** ***************************  ******

H. sapiens (IGLV1-40)      AGCCTGAGTGGTTC    299
M. putorius furo           AGCTTCAGTGCTCA    308
M. nigripes                AGCTTCAGTGCTCA    312
                           *** * **** *
```

**B**

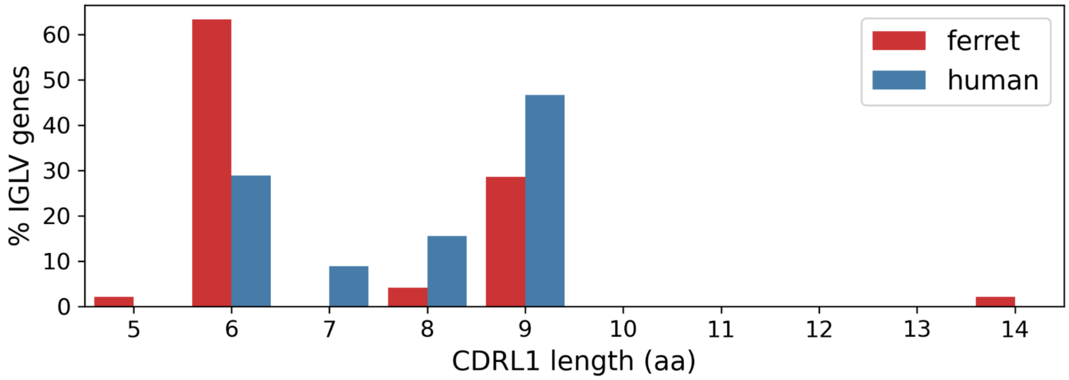


**Figure S1. The ferret long-CDRL1 IGLV gene. (A)** Alignment of the long-CDRL1 IGLV gene in the domestic ferret (*Mustela putorius furo*, middle line) against the closest human IGLV gene (IGLV1-40*01, top line) and the long-CDRL1 pseudogene in the black-footed ferret (*M. nigripes*, bottom line). **(B)** Amino acid lengths of CDRL1s across the IGLV genes of the domestic ferret (red) and human (blue). CDRL1s were computed using the IMGT definition (Lefranc, 2003).

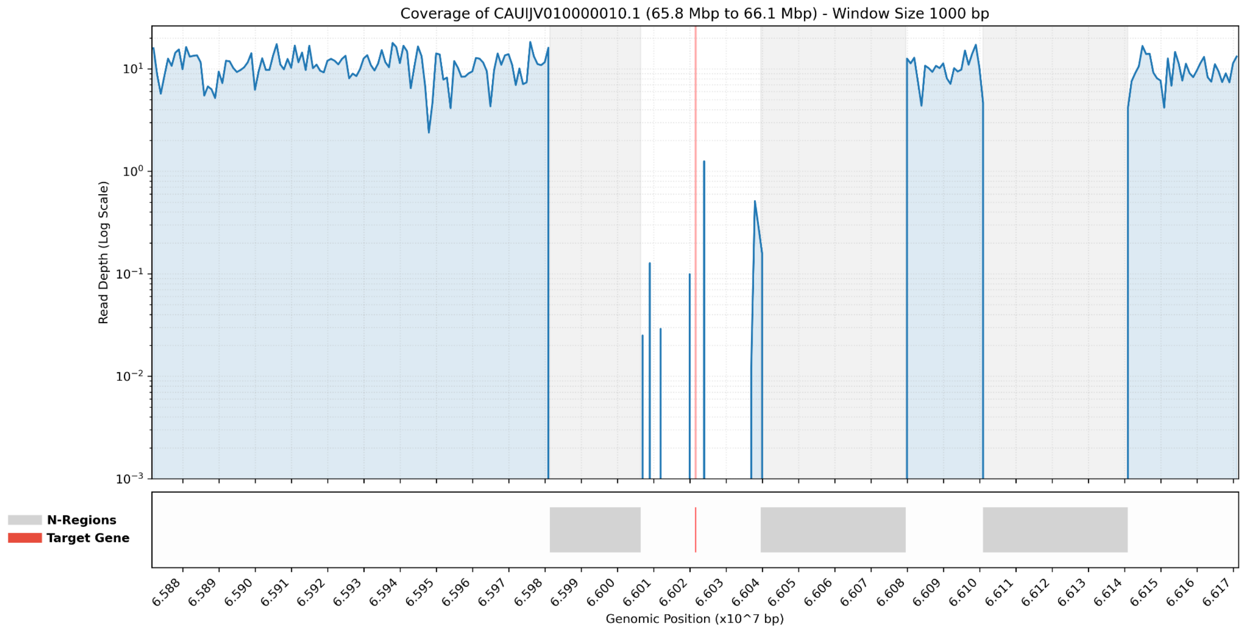


**Figure S2. Read depth of the steppe polecat subject from Inner Mongolia, China (ERR7198277) aligned to the genome of the steppe polecat subject from Altai, Russia (GCA_963422785.1).** The red line denotes the long-CDRL1 IGLV gene. Its position falls outside the N-regions (shown in gray) and aligns with a sharp drop in coverage indicating a deletion of the gene.

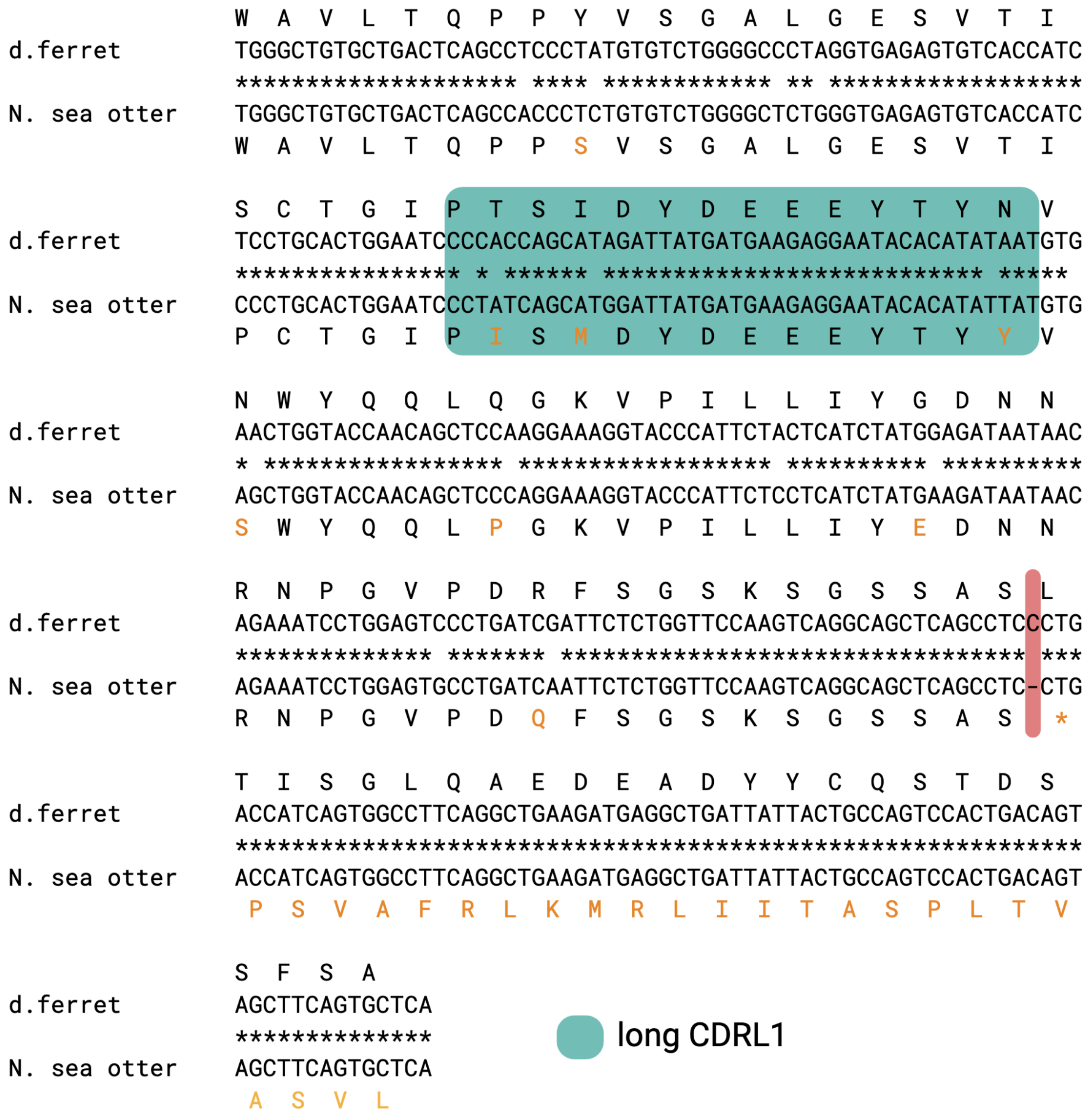


**Figure S3. Alignment of long-CDRL1 IGLV genes found in the domestic ferret (top line) and the Northern sea otter (bottom line).** Amino acids are shown above and below the first positions of corresponding codons. Amino acids in the Northern sea otter IGLV gene that differ from corresponding amino acids in the domestic ferret IGLV gene are shown in orange. The long-CDRL1 sequence is shown in green. The frameshifting one-nucleotide deletion in the Northern sea otter IGLV gene is shown in red.

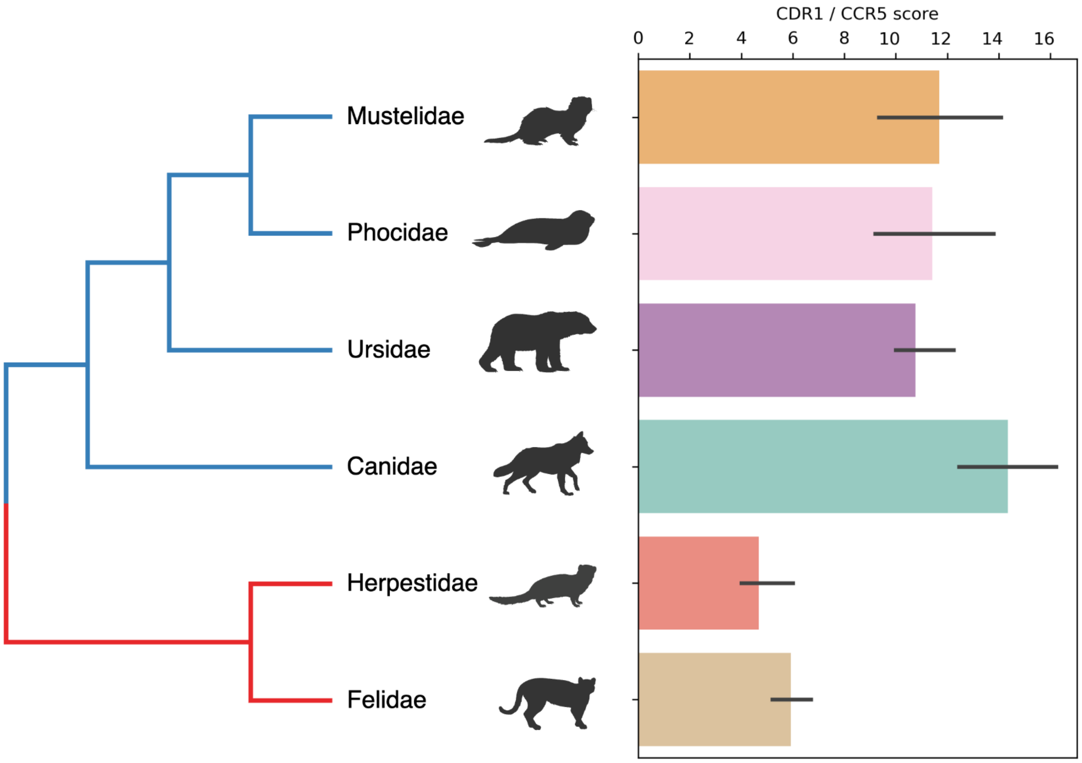


**Figure S4. The distribution of similarity scores between CDRL1s and CCR5s across six Carnivora families: Mustelidae, Phocidae, Ursidae, Canidae, Herpestidae, Felidae.** Caniformia and Feliformia families are shown by blue and red branches, respectively.

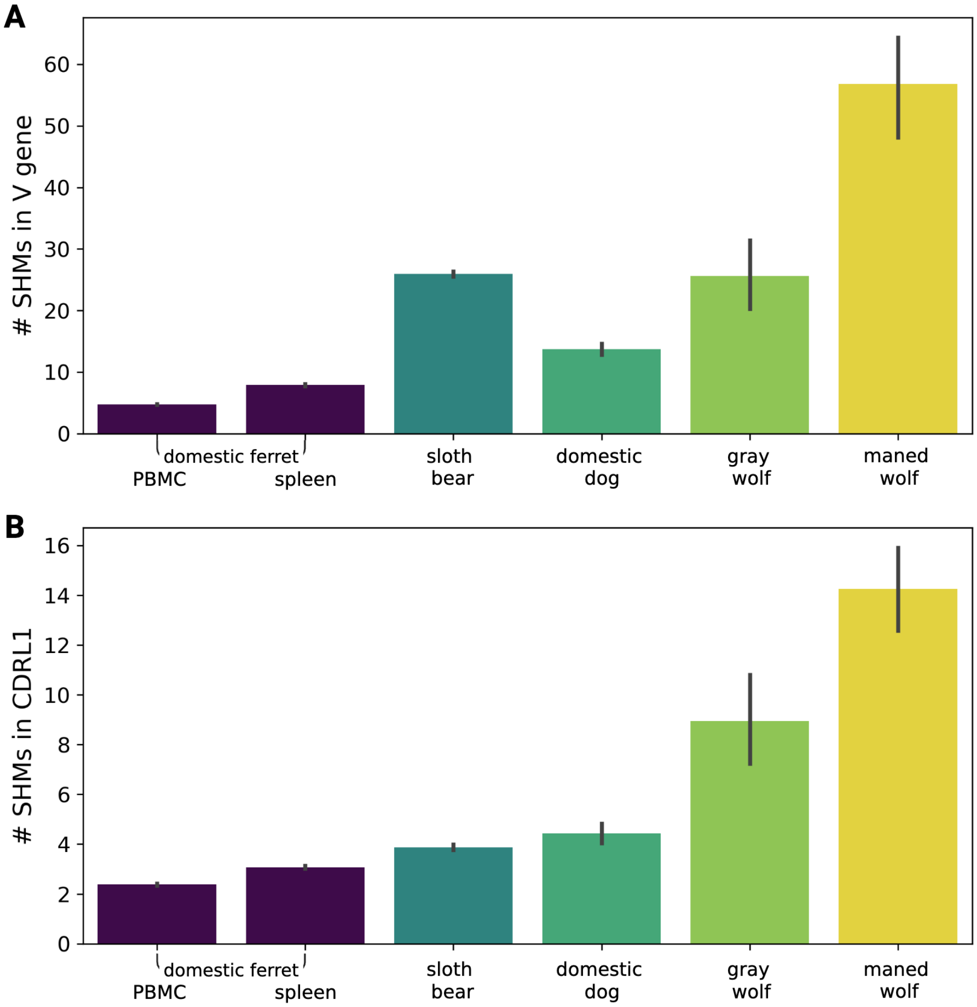


**Figure S5. Counts of somatic hypermutations (SHMs) in the V gene (A) and CDRL1 (B) of the long-CDRL1 IGL sequences across five carnivoran species: domestic ferret (PBMC and spleen), sloth bear (spleen), domestic dog (PBMC), gray wolf (spleen), and maned wolf (spleen).**

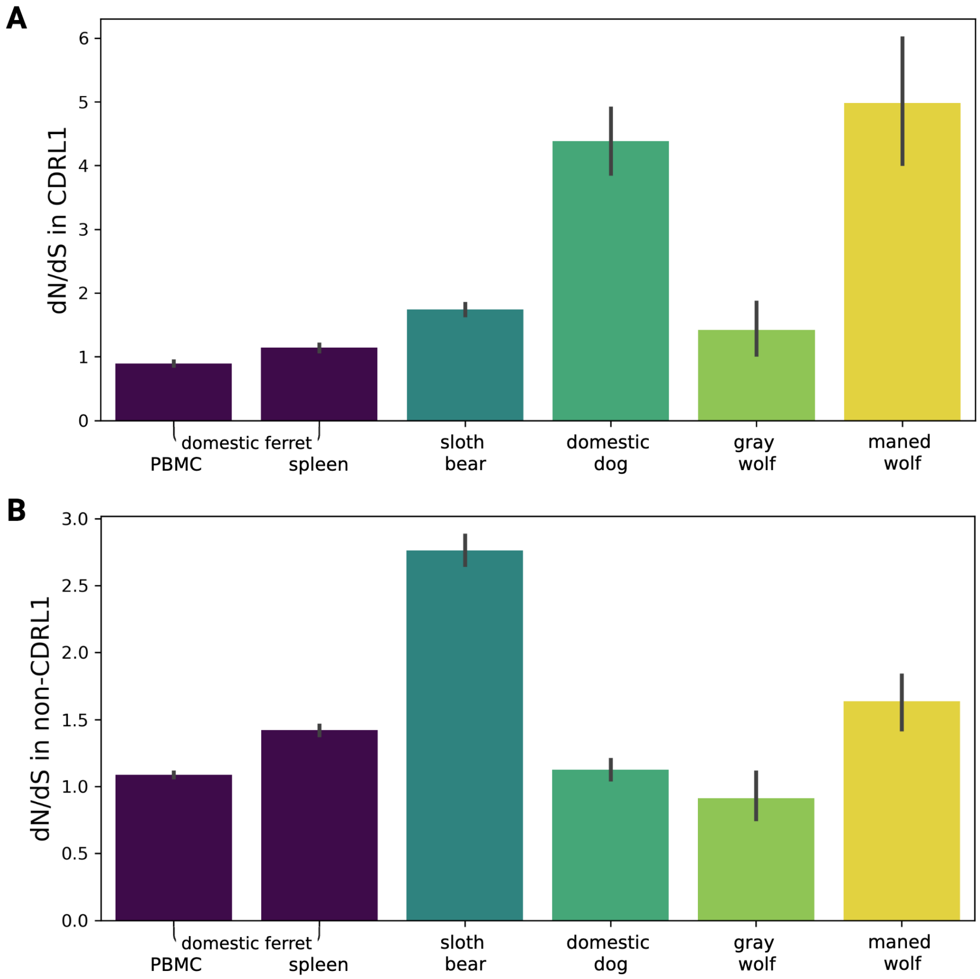


**Figure S6. dN/dS values computed based on somatic hypermutations in the CDRL1 (A) and non-CDRL1 parts (B) of the long-CDRL1 IGL sequences across five carnivoran species: domestic ferret (PBMC and spleen), sloth bear (spleen), domestic dog (PBMC), gray wolf (spleen), and maned wolf (spleen).**

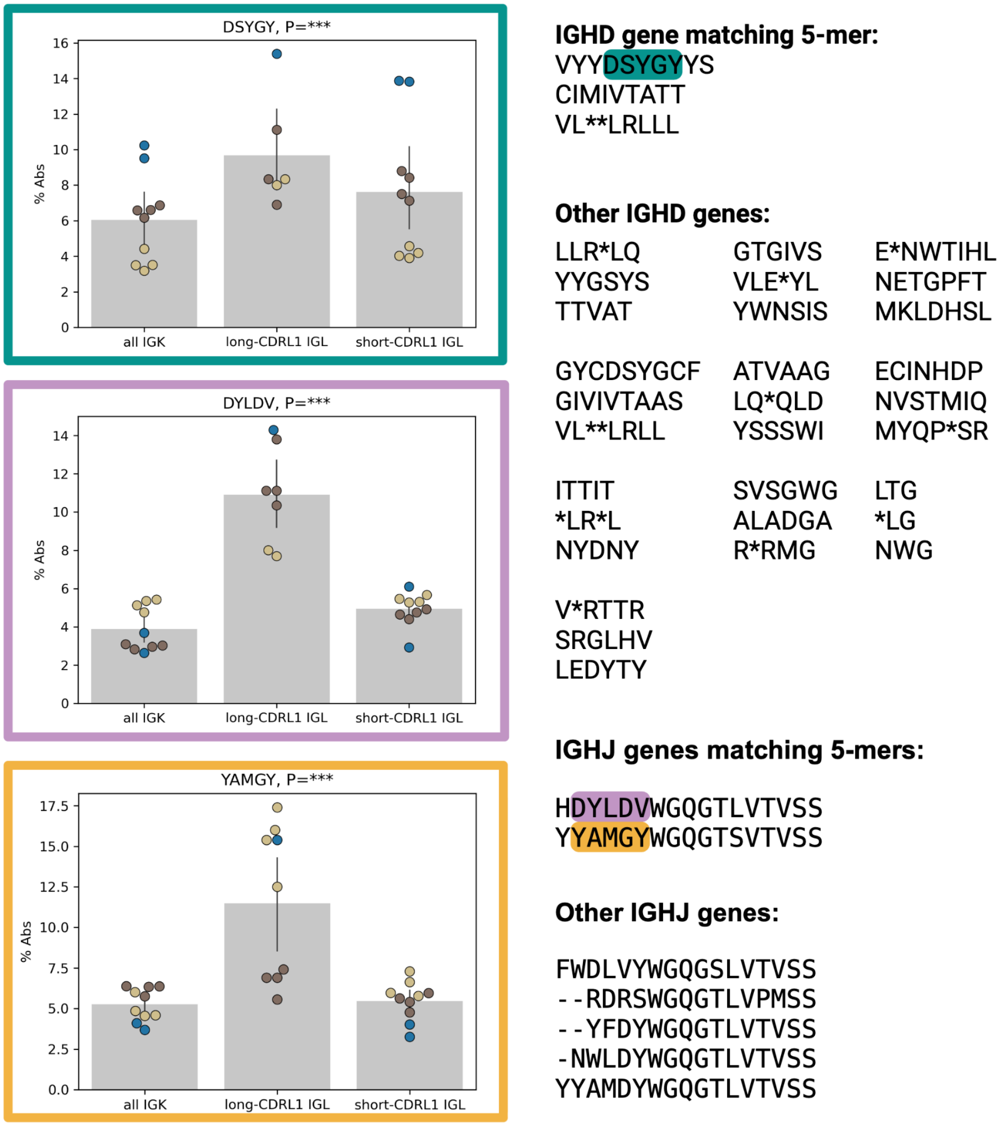


**Figure S7. 5-mers DSYGY (highlighted in green), DYLDV (highlighted in purple), and YAMGY (yellow) with significantly higher fractions in long-CDRL1 IGL Abs have perfect matches to one IGHD gene and two IGHJ genes of domestic ferret.** 10 other domestic ferret IGHD genes (three open reading frames per gene) and five other domestic ferret IGHJ genes (one open reading frame per gene) that do not have perfect matches to the identified 5-mers are shown on the right as well.

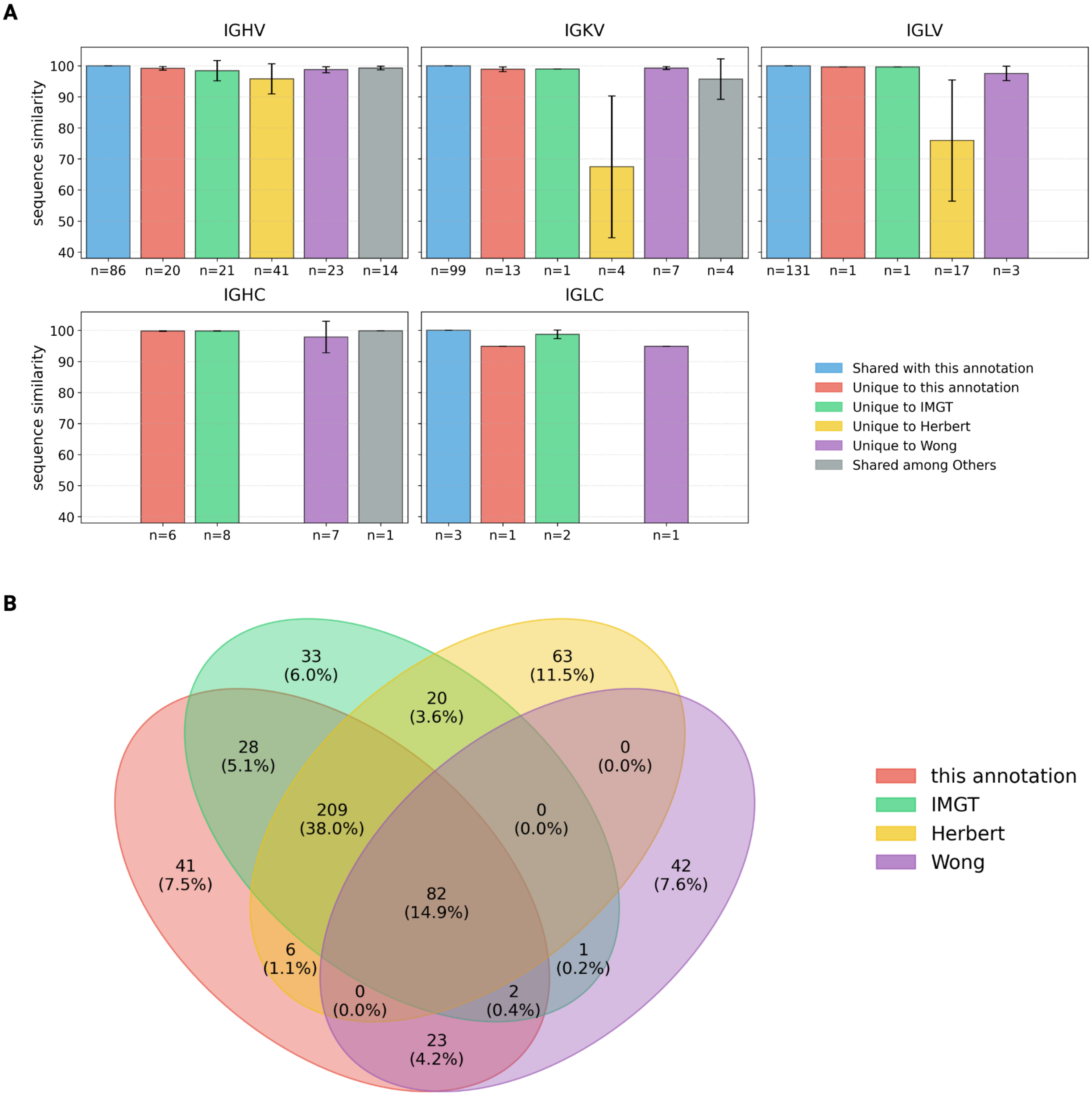


**Figure S8. Comparison of domestic ferret IG genes with existing IG annotations. (A)** Bar plots depict the sequence similarity for V and C genes with respect to corresponding genes from other databases across IGH, IGK and IGL loci. The counts of sequences (shown as n-values on the x-axes) are listed for each category and the error bar denotes the standard deviation. **(B)** The Venn diagram depicts counts and percentages of sequences shared across all four analyzed ferret IG gene annotations. Only perfect matches were visualized.

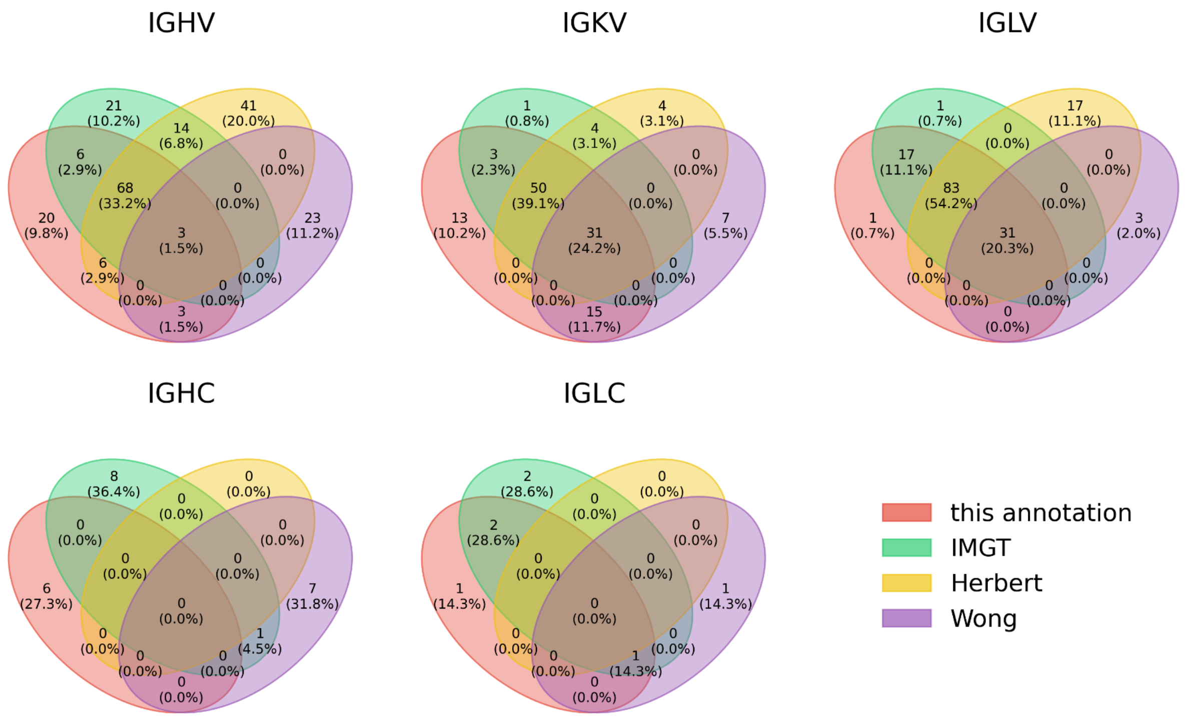


**Figure S9. Venn diagrams showing IGHV, IGKV, IGLV, IGHC, and IGLC genes shared among four analyzed annotations of domestic ferrets.** Only perfect matches were visualized.

### Supplemental Tables

| Species (common name) | Species (latin name) | Accession number | Sample location |
|---|---|---|---|
| Western polecat (N=7) | *Mustela putorius* | SRR30238149 | Pskov, Russia |
| | | ERR7256386 | Spain |
| | | ERR7256388 | Austria |
| | | ERR7256391 | France |
| | | ERR7256393 | Italy |
| | | ERR7256398 | Germany |
| | | ERR7256400 | United Kingdom |
| Steppe polecat (N=2) | *Mustela eversmanni* | GCA_963422785.1 | Altai Republic, Russia |
| | | ERR7198277 | Inner Mongolia, China |
| Black-footed ferret (N=8) | *Mustela nigripes* | SRR35734985 | Arizona, USA |
| | | SRR35734986 | |
| | | SRR35734987 | Virginia, USA (captive bred subjects) |
| | | SRR35734990 | |
| | | SRR35734988 | Colorado, USA (captive bred subjects) |
| | | SRR35734989 | |
| | | SRR35734991 | |
| | | SRR35734992 | |
| Back-striped weasel (N=1) | *Mustela strigidorsa* | GCA_048197275.1 | Vietnam |
| Siberian weasel (N=2) | *Mustela sibirica* | GCA_048197585.1 | Sakha Republic, Russia |
| | | SRR30238152 | Sikhote-Alin Nature Reserve, Russia |
| European mink (N=2) | *Mustela lutreola* | GCA_030435805.1 | Germany |
| | | GCA_964264385.1 | France |
| Least weasel (N=4) | *Mustela nivalis* | SRR30238153 | Sikhote-Alin Nature Reserve, Russia |
| | | SRR30238154 | Novosibirsk, Russia |
| | | SRR30238155 | Germany |
| | | GCA_019141155.1 | Poland |
| Stoat (N=2) | *Mustela erminea* | SRR30238150 | Republic of Sakha, Russia |
| | | SRR30238151 | |
| American mink (N=2) | *Neogale vison* | SRR33699353 | Louisiana, USA |
| | | SRR33699361 | Florida, USA |

**Table S1. Population data of Mustelinae species.**

| Species (eng) | Species (lat) | NCBI accession number |
|---|---|---|
| **Caniformia, Canidae** | | |
| Red fox | *Vulpes vulpes* | GCA_003160815.1 |
| Dingo | *Canis lupus dingo* | GCA_003254725.1 |
| German shepherd dog | *Canis lupus familiaris* | GCA_008641055.3 |
| African hunting dog | *Lycaon pictus* | GCA_001887905.1 |
| Bat-eared fox | *Otocyon megalotis* | GCA_017311455.1 |
| Santa Catalina Island fox | *Urocyon littoralis catalinae* | www.dnazoo.org/assemblies/urocyon_littoralis_catalinae |
| Greenland wolf | *Canis lupus orion* | GCA_905319855.2 |
| **Caniformia, Ursidae** | | |
| Spectacled bear | *Tremarctos ornatus* | www.dnazoo.org/assemblies/tremarctos_ornatus |
| American black bear | *Ursus americanus* | GCA_024610735.1 |
| Brown bear | *Ursus arctos* | GCA_023065955.1 |
| **Caniformia, Phocidae** | | |
| Northern elephant seal | *Mirounga angustirostris* | GCA_029215635.1 |
| Weddell seal | *Leptonychotes weddellii* | GCA_000349705.1 |
| Hawaiian monk seal | *Neomonachus schauinslandi* | GCA_002201575.2 |
| Spotted seal | *Phoca largha* | www.dnazoo.org/assemblies/phoca_largha |
| Harbor seal | *Phoca vitulina* | GCA_004348235.1 |
| **Caniformia, Otariidae** | | |
| California sea lion | *Zalophus californianus* | GCA_009762305.2 |
| **Caniformia, Mustelidae** | | |
| Northern sea otter | *Enhydra lutris kenyoni* | GCA_002288905.2 |
| Eurasian river otter | *Lutra lutra* | GCA_902655055.2 |
| Northern American river otter | *Lontra canadensis* | GCA_010015895.1 |
| Eurasian badger | *Meles meles* | GCA_922984935.2 |
| Stoat | *Mustela erminea* | GCA_009829155.1 |
| Stone marten | *Martes foina* | www.dnazoo.org/assemblies/martes_foina |
| European pine marten | *Martes martes* | GCA_963455335.1 |
| European mink | *Mustela lutreola* | GCA_030435805.1 |
| Least weasel | *Mustela nivalis* | GCA_964662115.1 |
| Sable | *Martes zibellina* | GCA_040938815.1 |
| **Caniformia, Mephitidae** | | |
| Plains spotted skunk | *Spilogale interrupta* | GCA_023159085.1 |
| **Feliformia, Herpestidae** | | |
| Dwarf mongoose | *Helogale parvula* | GCA_004023845.1 |
| Small Indian mongoose | *Urva auropunctata* | www.dnazoo.org/assemblies/urva_auropunctata |
| Meerkat | *Suricata suricatta* | GCA_006229205.1 |
| **Feliformia, Eupleridae** | | |
| Fossa | *Cryptoprocta ferox* | GCA_004023885.1 |
| **Feliformia, Felidae** | | |
| Clouded leopard | *Neofelis nebulosa* | GCA_028018385.1 |
| Canada lynx | *Lynx canadensis* | GCA_007474595.2 |
| Bobcat | *Lynx rufus* | GCF_022079265.1 |
| Domestic cat | *Felis catus* | GCA_018350175.1 |
| Mountain lion | *Puma concolor* | GCA_028749985.3 |
| Jaguarundi | *Herpailurus yagouaroundi* | GCA_014898765.1 |
| Jaguar | *Panthera onca* | GCA_004023805.1 |
| Amur tiger | *Panthera tigris altaica* | GCA_000464555.1 |
| Snow leopard | *Panthera uncia* | GCA_023721935.1 |

**Table S2. Publicly available genomes of carnivoran species selected for the comparative analysis.**

| Species | Latin name | AZA/SB# | Sex | Collection date | WGS tissue | Iso-Seq tissue |
|---|---|---|---|---|---|---|
| Black-footed ferret | *Mustela nigripes* | 7817 | M | 6-Nov-20 | lung | spleen |
| Cheetah | *Acinonyx jubatus* | 7221 | F | 27-Apr-22 | whole blood | - |
| Red panda | *Ailurus fulgens* | 213011 | M | 26-Apr-22 | spleen | spleen |
| Sloth bear | *Melursus ursinus* | 651 | M | 2-May-20 | heart | spleen |
| Gray wolf | *Canis lupus* | 274 | F | 2020 | spleen | spleen |
| Maned wolf | *Chrysocyon brachyurus* | 3153 | F | Jun-20 | spleen | spleen |

**Table S3. Information about samples of six newly sequenced carnivoran species: black-footed ferret, cheetah, gray wolf, maned wolf, red panda, and sloth bear.** The “AZA/SB#” column refers to Association of Zoos and Aquariums studbook numbers.

| Species | Haplotype 1 | | | Haplotype 2 | | |
|---|---|---|---|---|---|---|
| | Total genome size (Gbp) | N50 (Mbp) | L50 | Total genome size (Gbp) | N50 (Mbp) | L50 |
| Domestic ferret | 2.52 | 54.13 | 15 | 2.49 | 56.77 | 15 |
| Black-footed ferret | 2.23 | 50.00 | 12 | 2.36 | 60.60 | 12 |
| Cheetah | 2.40 | 49.45 | 16 | 2.40 | 40.45 | 14 |
| Red panda | 2.44 | 22.33 | 31 | 2.36 | 18.50 | 39 |
| Sloth bear | 2.39 | 18.79 | 39 | 2.18 | 0.26 | 2177 |
| Gray wolf | 2.43 | 35.67 | 23 | 2.45 | 36.88 | 25 |
| Maned wolf | 2.46 | 45.57 | 20 | 2.45 | 41.06 | 23 |

**Table S4. Characteristics of the new genome assemblies.**

| Subject | Tissue | # PE reads | # merged reads | # V(D)J reads | # IGL reads | # unique IGLs | # (%) long-CDRL1 IGLs |
|---|---|---|---|---|---|---|---|
| F17-22 | PBMC | 1,053,245 | 882,885 | 502,937 | 445,223 | 236,906 | 2563 (1.08%) |
| F17-22 | Spleen | 1,793,337 | 1,519,577 | 949,517 | 863,445 | 440,611 | 2820 (0.64%) |
| F63-22 | PBMC | 1,177,605 | 993,717 | 569,585 | 510,595 | 280,886 | 3823 (1.36%) |
| F63-22 | Spleen | 1,285,288 | 1,112,977 | 820,712 | 749,440 | 389,405 | 1884 (0.48%) |
| F64-22 | PBMC | 578,468 | 491,991 | 267,271 | 236,515 | 131,889 | 2375 (1.80%) |
| F64-22 | Spleen | 1,835,446 | 1,589,997 | 1,133,012 | 1,034,017 | 493,584 | 4204 (0.85%) |

**Table S5. Characteristics of the new domestic ferret Rep-seq IGL libraries.**

| Species | # Iso-Seq reads | # (%) IG reads | # (%) IGH reads | # (%) IGK reads | # (%) IGL reads | # (%) long-CDRL1 IGLs |
|---|---|---|---|---|---|---|
| Black-footed ferret (*Mustela nigripes*) | 4,204,992 | 94,737 (2.25%) | 48,535 (51.23%) | 23,619 (24.93%) | 22,583 (23.84%) | 0 (0.00%) |
| Red panda (*Ailurus fulgens*) | 3,659,921 | 15,269 (0.42%) | 7593 (49.73%) | 5148 (16.56%) | 2528 (33.71%) | 19 (1.76%) |
| Sloth bear (*Melursus ursinus*) | 4,829,413 | 926,740 (19.19%) | 495,321 (53.45%) | 61,199 (6.60%) | 370,220 (39.95%) | 1837 (2.88%) |
| Gray wolf (*Canis lupus*) | 4,418,571 | 198,850 (4.50%) | 131,704 (66.23%) | 10,379 (5.22%) | 56,767 (28.55%) | 68 (0.36%) |
| Maned wolf (*Chrysocyon brachyurus*) | 4,102,039 | 448,072 (10.92%) | 252,076 (56.26%) | 11,513 (2.57%) | 184,483 (41.17%) | 77 (0.04%) |

**Table S6. Characteristics of the new Iso-Seq libraries for the black-footed ferret, red panda, sloth bear, gray wolf, and maned wolf.**

| Subject | # PE reads | # merged reads | # V(D)J reads | # IGL reads | # unique IGLs | # (%) long-CDRL1 IGLs |
|---|---|---|---|---|---|---|
| **PBMC of the domestic ferrets, project PRJNA1256357** | | | | | | |
| SRR33336524 | 667,082 | 614,565 | 458,274 | 377,795 | 209,524 | 9335 (4.46%) |
| SRR33336525 | 1,292,860 | 1,161,166 | 1,050,948 | 843,149 | 578,273 | 18,224 (3.15%) |
| **PBMC of the domestic dogs, project PRJNA790470** | | | | | | |
| SRR17271941 (Golden Retriever) | 3,176,721 | 519,516 | 132,897 | 104,671 | 45,284 | 102 (0.23%) |
| SRR17271942 (Labrador Retriever) | 2,491,353 | 328,710 | 95,960 | 87,149 | 39,564 | 68 (0.17%) |
| SRR17271945 (Newfoundland) | 3,291,083 | 170,821 | 38,460 | 33,497 | 20,483 | 42 (0.21%) |
| SRR17271967 (German Shepherd) | 863,980 | 236,524 | 74,273 | 68,287 | 26,695 | 9 (0.03%) |
| SRR17271977 (Catahoula Hound) | 7,265,709 | 699,168 | 148,549 | 127,077 | 48,116 | 216 (0.45%) |
| SRR17271989 (Labrador Retriever) | 720,440 | 180,512 | 60,248 | 55,697 | 22,167 | 44 (0.20%) |
| SRR17271990 (Labrador Retriever) | 2,040,076 | 505,454 | 118,997 | 108,097 | 40,170 | 20 (0.05%) |

**Table S7. Characteristics of publicly available Rep-Seq datasets of the domestic ferrets and domestic dogs used in the study.**

| Subject | Run | # complete Abs (one HC + one LC) | # (%) IGL Abs | # (%) long-CDRL1 IGL Abs |
|---|---|---|---|---|
| S2, Splenocytes | SRR24488356 | 1294 | 696 (53.79%) | 25 (3.59%) |
| | SRR24488357 | 1271 | 705 (55.47%) | 26 (3.69%) |
| | SRR24488368 | 1258 | 669 (53.18%) | 24 (3.59%) |
| | SRR24488369 | 1236 | 664 (53.72%) | 23 (3.46%) |
| S3, PBMC | SRR24488362 | 1776 | 829 (46.68%) | 29 (3.50%) |
| | SRR24488363 | 1812 | 855 (47.19%) | 36 (4.21%) |
| | SRR24488370 | 1762 | 823 (46.71%) | 27 (3.28%) |
| | SRR24488371 | 1729 | 802 (46.39%) | 29 (3.62%) |

**Table S8. Characteristics of publicly available scRNA-Seq data of the domestic ferrets used in the study.**
